# Exploring high-quality microbial genomes by assembling short-reads with long-range connectivity

**DOI:** 10.1101/2022.09.07.506963

**Authors:** Zhenmiao Zhang, Jin Xiao, Hongbo Wang, Chao Yang, Yufen Huang, Zhen Yue, Yang Chen, Lijuan Han, Kejing Yin, Aiping Lyu, Xiaodong Fang, Lu Zhang

## Abstract

Despite long-read sequencing enables to generate complete genomes of unculturable microbes, its high cost hinders its widespread application in large cohorts. An alternative method is to assemble short-reads with long-range connectivity, which can be a cost-effective way to generate high-quality microbial genomes. We developed Pangaea to improve metagenome assembly using short-reads with physical or virtual barcodes. It adopts a deep-learning-based binning algorithm to assemble the co-barcoded reads with similar sequence contexts and abundances to improve assemblies of high- and medium-abundance microbes. Pangaea also leverages a multi-thresholding reassembly strategy to refine assembly for low-abundance microbes. We benchmarked Pangaea with linked-reads and a combination of short- and long-reads from mock communities and human gut metagenomes. Pangaea achieved significantly higher contig continuity as well as more near-complete metagenome-assembled genomes (NCMAGs) than the existing assemblers. Pangaea was also observed to generate three complete and circular NCMAGs on the human gut microbiomes.

## Introduction

Metagenome assembly is one of the main steps to reconstruct microbial genomes from culture-free metagenomic sequencing data [1]. Cost-effective short-read sequencing technologies have been widely applied to generate high-quality microbial reference genomes from large cohorts of human gut microbiomes [2, 3, 4]. However, the short read length (100bps-300bps) may not allow us to resolve intra-species repetitive regions and inter-species conserved regions [5] or to achieve complete microbial genomes. The emerging long-read sequencing technologies such as PacBio continuous long-read sequencing (PacBio CLR) [6], Oxford Nanopore sequencing (ONT) [7] and PacBio HiFi sequencing [8], have shown their superiority to short-read sequencing in generating metagenome-assembled genomes (MAGs) with high continuity or producing complete and circular MAGs using the long-range connectivity they provided [9, 10, 11]. Despite potential benefits, the high cost of long-read sequencing makes deep sequencing impracticable and continues to hinder its application in population-scale or clinical studies [12]. In our previous research [13], long-reads generated fewer high-quality MAGs than short-reads due to insufficient sequencing depth. As an alternative way to deep long-read sequencing, some studies [14, 15, 16, 13] suggested utilizing cost-effective short-reads with long-range connectivity for metagenome assembly. The long-range connectivity could be derived from physical barcodes (e.g., linked-reads) or virtual links by other long-fragment sequencing technologies (e.g. long-reads).

Linked-read sequencing attaches identical barcodes (physical barcodes) to the short-reads if they are derived from the same long DNA fragment. Before the discontinuation, 10x Chromium was the most widely used linked-read sequencing technology, generating contigs with high continuity and producing more near-complete metagenome-assembled genomes (NCMAGs; **Methods**) than short-read sequencing [13, 14]. However, co-barcoded short-reads of 10x have a high chance of being derived from multiple DNA fragments (average number of fragments per barcode [*N_F/B_*] is 16.61; **Methods** and **Supplementary Note 1**), which may complicate the deconvolution of complex microbial communities. Recently, MGI and Universal Sequencing Technology released their linked-read sequencing technologies, namely single-tube Long Fragment Read (stLFR) [17] and Transposase Enzyme-Linked Long-read Sequencing (TELL-Seq) [18]. The barcoding reactions of these technologies occur on billions of microbeads, leading to much higher barcode specificity (*N_F/B_* = 1.54 for stLFR; *N_F/B_* = 4.26 for TELL-Seq; **Supplementary Note 1**). The co-barcoded short-reads of these technologies are more likely to come from the same genomic regions.

Several assemblers have been developed for assembling linked-reads: (i) Athena [14], which fills the gaps between contigs by recruiting the 10x co-barcoded reads for local assembly; (ii) cloudSPAdes [19], which reconstructs the long DNA fragments in the assembly graph for solving the shortest superstring problem to improve contig continuity; (iii) Supernova [20], which was developed for diploid human genome assembly by allowing two paths in megabubble structures based on a series of modification of assembly graph using 10x co-barcoded reads; (iv) MetaTrass [21], which groups the stLFR co-barcoded reads by reference-based taxonomic annotation and applies Supernova to assemble the genome of each identified species. With the exception of MetaTrass, all the other three tools were developed for 10x linked-reads with low barcode specificity. We exclude MetaTrass for comparison in this paper because it relies on the available microbial reference genomes and thus has a limited capability to discover novel species. There is still a lack of efficient tools that could fully exploit the long-range connectivity of short-reads from high-specificity barcodes to improve *de novo* metagenome assembly.

Long-range connectivity could also be provided by other long-fragment sequencing technologies, which can be used in conjunction with short-reads for hybrid assembly. The hybrid assembly is typically performed by combining high-depth short-reads with shallow-depth long-reads. This approach takes advantage of the high base quality of short-reads for contig assembly and the long-range connectivity of long-reads for contig extension. Two hybrid assemblers were developed for metagenome assembly: (i) hybridSPAdes [16] maps long-reads to the assembly graph from short-reads and utilizes the long-range connectivity to resolve uneven path depth and repetitive sequences in the graph; (ii) OPERA-MS [15] aligns long-reads to the contigs assembled from short-reads to construct a scaffold graph and groups contigs based on microbial reference genomes followed by gap filling in each cluster.

Previous studies showed that read subsampling was an effective strategy for assembling large complex metagenomic datasets [22]. It could improve the assembly of high-abundance microbes [23], but result in poor quality in assembling low-abundance microbes [24] due to insufficient reads. In a metagenome community, microbes may have highly uneven abundances [25]. Sequences obtained from low-abundance microbes can result in spurious edges with shallow depths in the assembly graph. Distinguishing these connections from edges originating from sequencing errors becomes a particularly challenging task [26]. We introduce Pangaea, a metagenome assembler, to improve metagenome assembly with three modules. Firstly, Pangaea applies co-barcoded short-read clustering and assembly to reduce the complexity of metagenomic sequencing data. The read binning strategy has been proven advantageous in metagenome assembly [27, 28, 29]. It could be a more sophisticated read downsampling strategy to improve the assemblies of high- and medium-abundance microbes. However, the existing tools are impractical when it comes to handling millions of short-reads within acceptable time and memory limitations. Pangaea groups co-barcoded short-reads rather than grouping independent short-reads, as the co-barcoded reads have highly specific barcodes (**Methods**). Short-reads that belong to the same cluster have lower complexity than the original dataset and are assembled separately. Secondly, Pangaea adopts a multi-thresholding reassembly step to refine the assembly of low-abundance microbes using different abundance thresholds to handle the uneven abundances of microbes (**Methods**). The data from high-abundance microbes are gradually removed from the assembly graph, thus the sequences from various levels of low-abundance microbes are preserved. Athena [14] demonstrated that enhancing contig continuity can be achievable by combining the original short-read assembly with contig local assembly based on co-barcoded reads. Thirdly, Pangaea integrates the assemblies from the above two modules, original short-read assembly and local assembly to improve contig continuity (Methods). We benchmarked Pangaea using linked-reads with physical barcodes from linked-read sequencing of the mock and human gut metagenomes, and evaluated its generalizability using short-reads with virtual barcodes generated from their alignments on long-reads (**Methods**). For linked-reads, we compared Pangaea with two short-read assemblers (metaSPAdes [30] and MEGAHIT [31]) and three linked-read assemblers (cloudSPAdes, Supernova and Athena). We found Pangaea achieved substantially better contig continuity and more NCMAGs than the other tools on all datasets. It also generated three complete and circular microbial genomes for the three real complex microbial communities. For short-reads with virtual barcodes, Pangaea could substantially improve contig continuity and generate better assemblies for both high- and low-abundance microbes than the short-read and hybrid assemblers.

## Results

### Workflow of Pangaea

Pangaea is a *de novo* metagenome assembler designed for short-reads with long-range connectivities represented by their attached barcodes (**Figure 1 a**). The barcodes can be generated physically (from linked-read sequencing with high barcode specificity, e.g., stLFR and TELL-Seq) or virtually (from long-reads; **Methods**). The co-barcoded short-reads are considered to be derived from the same long DNA fragments. The core algorithms of Pangaea are designed to reduce the complexity of metagenomic sequencing data (for high- and medium-abundance microbes) and deal with the uneven abundances of involved microbes (for low-abundance microbes) based on the barcodes with high specificity. Pangaea contains three main modules: (i) Co-barcoded read binning. This module is intended to reduce metagenomic complexity before assembly and is mainly used to improve the assembly of high- and medium-abundance microbes from real-world datasets with high complexity. Pangaea collects the co-barcoded short-reads and represents them using *k*-mer histograms and TNFs (**Methods**) to overcome feature instability in grouping short-reads individually. These features enable Variational Autoencoder (VAE) to represent co-barcoded short-reads in low-dimensional latent space, where the latent variable follows a standard Gaussian distribution (**Methods**; **Supplementary Note 2**). Pangaea adopts a weighted sampling strategy in VAE training to balance the number of co-barcoded short-reads from microbes with different abundances (**Methods**). The *k*-mer histogram (*X_A_*) for each specific barcode can be modeled as the mixture of two Poisson distributions, one represents the erroneous *k*-mers with extremely low frequency [32], and the other represents the true *k*-mer abundances [33]. We observed there was a negative correlation between *max*(*X_A_*) and the abundance of the co-barcoded reads (**Supplementary Figure 1**). Co-barcoded reads from low-abundance microbes should have a higher chance of being sampled, thus we used *max*(*X_A_*)^2^ as the sampling weight of each barcode (**Methods**). Pangaea groups co-barcoded short-reads in the latent space using RPH-kmeans [34], which is beneficial if the bins have uneven sizes (**Methods**) and assembles the short-reads in each bin independently. (ii) Multi-thresholding reassembly. This module refines the assembly of low-abundance microbes by removing the sequences of high-abundance microbes from the assembly graph gradually (**Figure 1 b**; **Methods**). This module aligns short-reads to contigs and removes them if they are from the contigs with abundances above certain thresholds (**Methods**). The remaining reads are used to reassemble the contigs of low-abundance microbes. (iii) Ensemble assembly. This module is to eliminate the impact of mis-binning on final assembly results. Athena [14] shows the integration of original short-read assembly and local assembly can improve metagenome assembly using the overlaplayout-consensus (OLC) strategy. Following the same strategy, Pangaea additionally incorporates the assemblies from the previous two modules (**Methods**). Pangaea would finally integrate the ensemble assembly with the assembly from the available assembly tools for the short-reads with virtual barcode using quickmerge (**Methods**). This is optional for the assembly of short-reads with physical barcodes.

**Figure 1:**
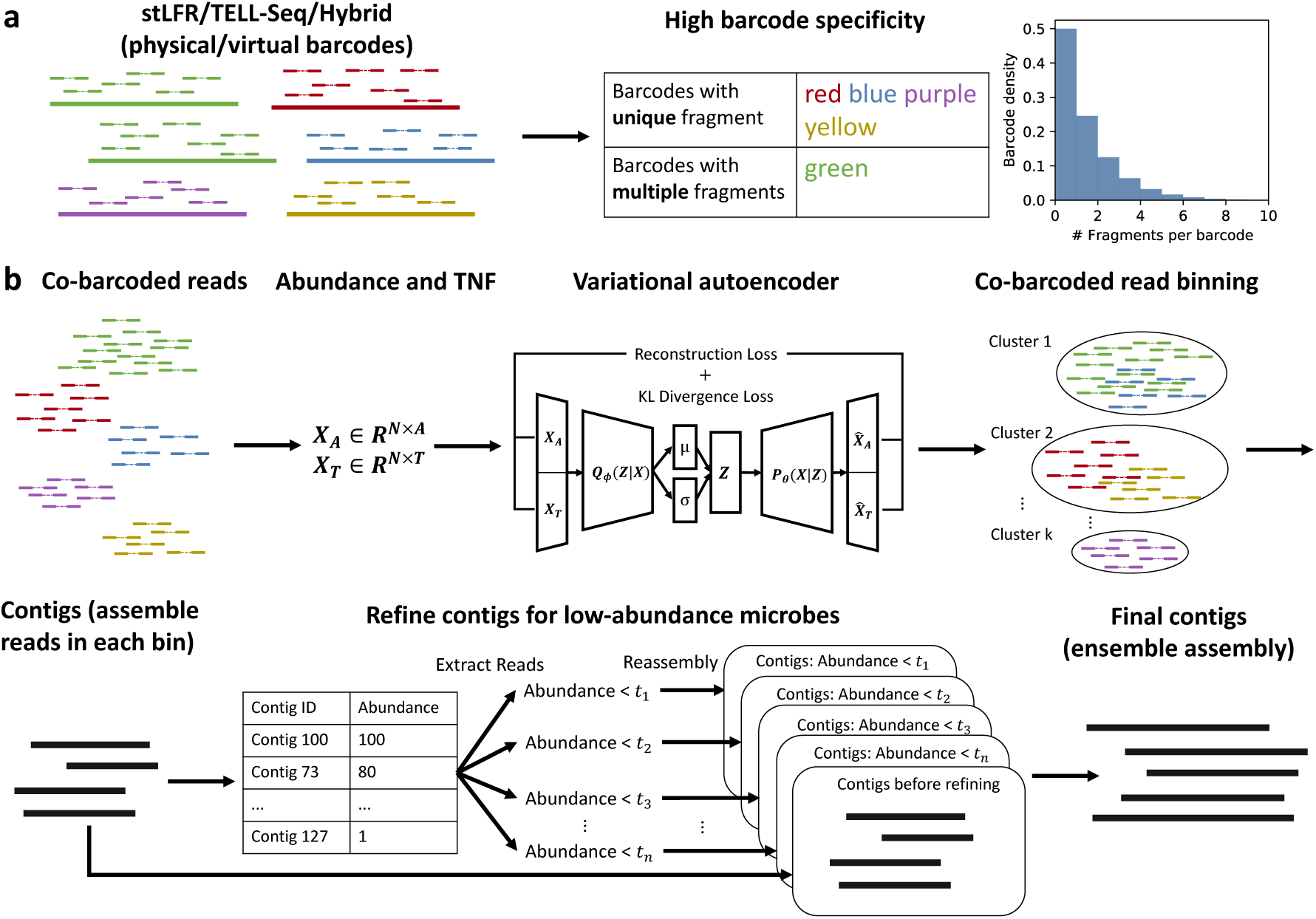
Workflow of Pangaea on stLFR and TELL-Seq linked-reads (physical barcodes) and short-reads with virtual barcodes from long-reads. (**a**) High barcode specificity for stLFR and TELL-Seq linked-reads. (**b**) Pangaea extracts features including *k*-mer histograms and TNFs from co-barcoded reads. The features are concatenated and used to represent reads in low-dimensional latent space using a variational autoencoder. The embeddings of co-barcoded reads are clustered by RPH-kmeans. Pangaea assembles the reads from each bin independently and adopts a multi-thresholding reassembly strategy to improve the assemblies for low-abundance microbes. Ensemble assembly integrates the contigs from different strategies using OLC algorithm.

### Co-barcoded read binning increases NCMAGs from medium- and high-abundance microbes

Co-barcoded read binning in the first module is a core step of Pangaea. We demonstrate this module significantly improved the assembly for high- and medium-abundance microbes, particularly for the real complex microbial communities. We sequenced the stLFR (132.95Gb) and TELL-Seq (173.2Gb) linked-reads (**Methods**) of the ATCC-MSA-1003 mock community containing 20 strains (whose reference genomes are available; **Supplementary Table 1**) with loadings from 0.02% to 18% to evaluate co-barcoded read binning on the low-complexity mock dataset. We also prepared three real human gut microbiome datasets (S1, S2, and S3) with stLFR sequencing (**Supplementary Table 2**; **Methods**) to validate the algorithm on real complex microbial communities.

As there were no available binning tools for linked-reads, we compared the VAE-based binning algorithm in Pangaea with two existing long-read binning tools, METABCC-LR (v2.0.0) [33] and LRBinner (v2.1) [35] on ATCC-MSA-1003. In order to enable these long-read binning tools on linked-reads, the co-barcoded reads were connected using a single “N” into long-reads. LRBinner failed to finish the binning task within three weeks with 100 threads, so we excluded it for further comparison. The VAE-based binning algorithm achieved a higher overall F1 score and adjusted rand index (ARI) than METABCC-LR (Panagea: F1 = 0.6144, ARI = 0.3764; METABCC-LR: F1 = 0.5887; ARI = 0.1704; **Supplementary Table 3**). The VAE-based binning particularly improved the binning performance for low-abundance microbes (abundance: 0.02%), where it achieved an F1 score of 0.6073 and ARI of 0.4624. The performance of METABCC-LR was almost the same as random binning (**Supplementary Table 3**).

We compared the metagenome assemblies of Panagea with (*ASM_B_*) and without (*ASM_¬B_*) read binning to investigate the impact of co-barcoded read binning on final assembly results. For TELL-Seq of ATCC-MSA-1003, *ASM_B_*had a higher overall NA50 (649.67Kb) than *ASM_¬B_* (601.67Kb; **Supplementary Table 4**). Considering the individual strains, *ASM_B_* achieved higher NGA50s for 7 out of the 10 strains with abundances higher than 1% (**Supplementary Table 4**). For the human gut microbiomes, *ASM_B_* obtained comparable N50s and more NCMAGs than *ASM_¬B_* on the three samples (S1: *ASM_B_* = 24, *ASM_¬B_* = 20; S2: *ASM_B_* = 17, *ASM_¬B_* = 11; S3: *ASM_B_* = 9, *ASM_¬B_* = 8). These NCMAGs generated by *ASM_B_*were commonly observed from high-abundance microbes (average depths of NCMAGs: S1 = 526.6X; S2 = 211.19X; S3 = 256.52X). Next, we evaluated the impact of bin number on the binning performance and final assembly of ATCC-MSA-1003. We observed a large bin number resulted in high binning precision and low recall (**Supplementary Figure 2**). The bin number is a trade-off between generating bins with low complexities (large bin number) or keeping more reads from the same microbes in the same bin (small bin number). Although the bin numbers would influence the read binning performance, the assembly results were robust if bin numbers varied in a certain range (e.g. 10-20 in ATCC-MSA-1003 with 20 strains; **Supplementary Figure 3**). In general, datasets with high complexity should be assigned a large bin number. We used 15 and 30 as bin numbers for the datasets from ATCC-MSA-1003 and the human gut microbiomes, respectively.

### Multi-thresholding reassembly improves assemblies of low-abundance mi-crobes

Pangaea improves assemblies of low-abundance microbes by gradually removing the reads of high-abundance microbes from the assembly graph using multiple abundance thresholds (represented by T). We compared assemblies using different settings of T and found T = *{*10, 30*}* was the best choice (**Supplementary Note 3**), which was used in the subsequent experiments. We compared the assemblies with and without multi-thresholding reassembly and found this module could increase the NGA50s of 5 strains with abundances less than 1% (out of 6 strains, covered by more 50% genomes; **Supplementary Table 5**). It could generate more sequences from the long contigs (*>*10Kb) of 6 strains with abundances less than 1% (out of 7 strains with contigs longer than 10Kb; **Supplementary Table 5**). For the three human gut microbiomes, the assembly with multi-thresholding reassembly could also generate two more NCMAGs.

### Barcode specificity is critical for linked-read assembly

We sequenced ATCC-MSA-1003 using stLFR, TELL-Seq linked-read sequencing and collected 10x linked-reads [36] (**Supplementary Table 2**; **Methods**) to evaluate the influence of barcode specificity in metagenome assembly. Linked-reads of stLFR yielded the lowest *N_F/B_*(*N_F/B_*= 1.54), and TELL-Seq linked-reads yielded a slightly higher value (*N_F/B_* = 4.26). Both values were much lower than that obtained from 10x linked-reads (*N_F/B_*= 16.61; **Supplementary Note 1**).

The contigs of Pangaea from stLFR and TELL-Seq linked-reads had substantially higher N50 (22.42 times on average; Figure2 a) and higher overall NA50 (11.55 times on average; **Supplementary Table 6**) than the contigs of Athena and Supernova from 10x linked-reads. For the 15 strains with abundance higher than 0.1% (**Supplementary Table 7**), Pangaea on stLFR and TELL-Seq linked-reads also achieved significantly higher per-strain NA50 (**Figure 2 d**) and NGA50 (**Figure 2 g**) than the assemblies of Athena and Supernova using 10x linked-reads. For the remaining 5 strains with abundances of 0.02%, Pangaea on stLFR (average genome fraction: 33.49%) and TELL-Seq (average genome fraction: 23.55%) linked-reads obtained much higher genome fractions than Athena (average genome fraction: 3.99%) and Supernova (average genome fraction: 6.29%) on 10x linked-reads (**Supplementary Table 8**). These results suggest that short-reads with high barcode specificity could produce better metagenome assemblies using Pangaea.

**Figure 2:**
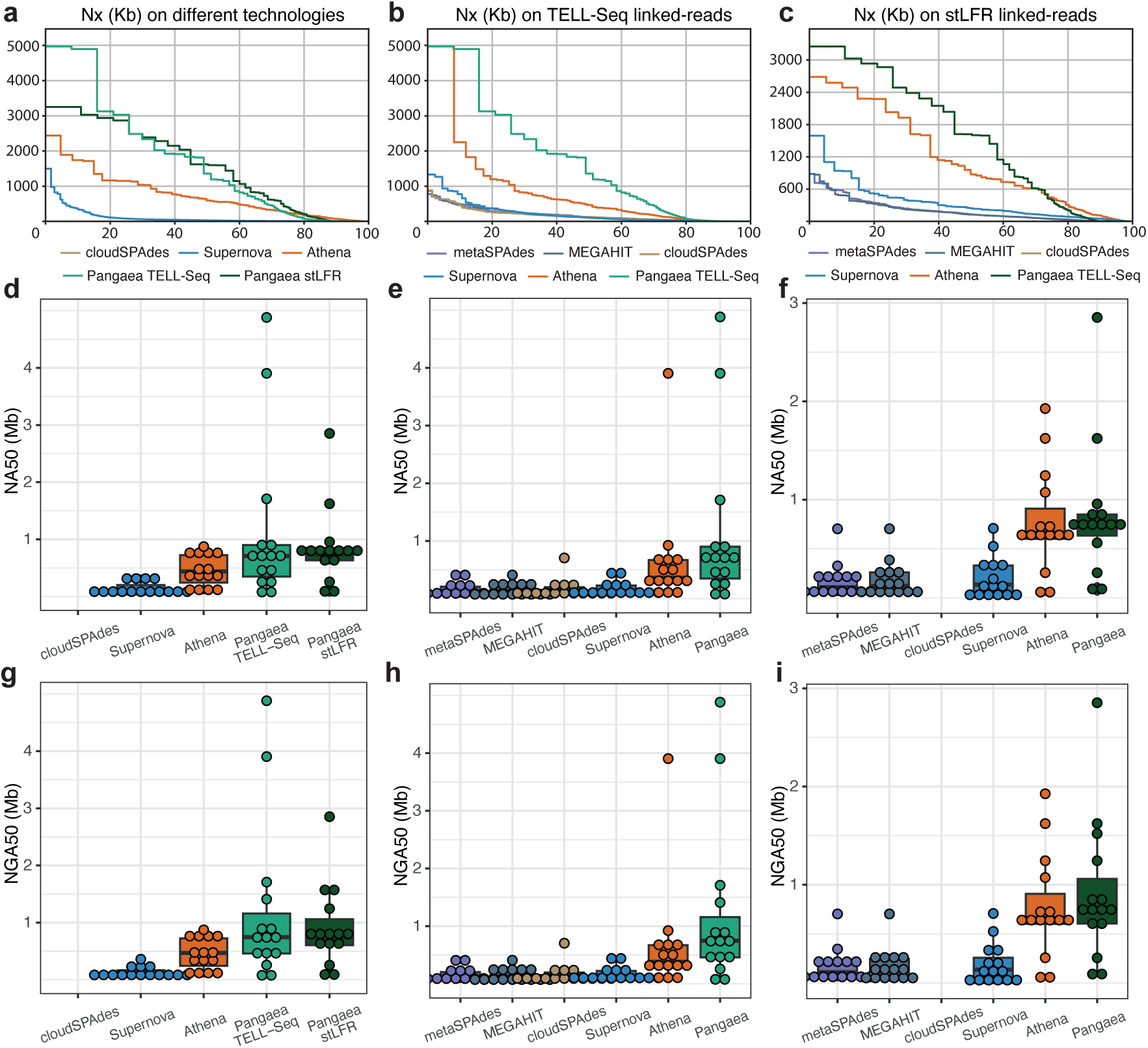
Nx, with x ranging from 0 to 100, for different linked-read sequencing technologies (**a**; except Panagea, all assemblers were applied to 10x linked-reads), TELL-Seq linked-reads (**b**), and stLFR linked-reads (**c**) from ATCC-MSA-1003 mock community. NA50 and NGA50 for the 15 strains with abundances higher than 0.1% assembled for different linked-read sequencing technologies (**d and g**; except Panagea, all assemblers were applied to 10x linked-reads), TELL-Seq linked-reads (**e and h**), and stLFR linked-reads (**f and i**) on ATCC-MSA-1003. cloudSPAdes was unavailable for 10x linked-reads and stLFR linked-reads, as it requires extremely large memory (*>*1TB) on these datasets.

### Pangaea generated high-quality assemblies for both high- and low-abundance microbes on mock linked-read datasets

We used two mock linked-read datasets (ATCC-MSA-1003 and ZYMO) to compare Pangaea to Athena, Supernova, cloudSPAdes, MEGAHIT, and metaSPAdes. For ATCC-MSA-1003, we benchmarked Pangaea on both TELL-Seq and stLFR linked-reads. For TELL-Seq (**Table 1**; **Figure 2 b**), Pangaea achieved the highest N50 (1,360.32Kb) and overall NA50 (649.47Kb) when compared with the statistics achieved by Athena (N50: 466.50Kb; NA50: 361.57Kb), Supernova (N50: 102.76Kb; NA50: 97.31Kb), cloudSPAdes (N50: 127.42Kb; NA50: 118.16Kb), MEGAHIT (N50: 128.07Kb; NA50: 112.51Kb) and metaSPAdes (N50: 112.34Kb; NA50: 105.63Kb). The contig N50 and overall NA50 for Pangaea on stLFR linked-reads were akin to the results on TELL-Seq linked-reads (**Table 1**; **Figure 2 c**).

**Table 1:**
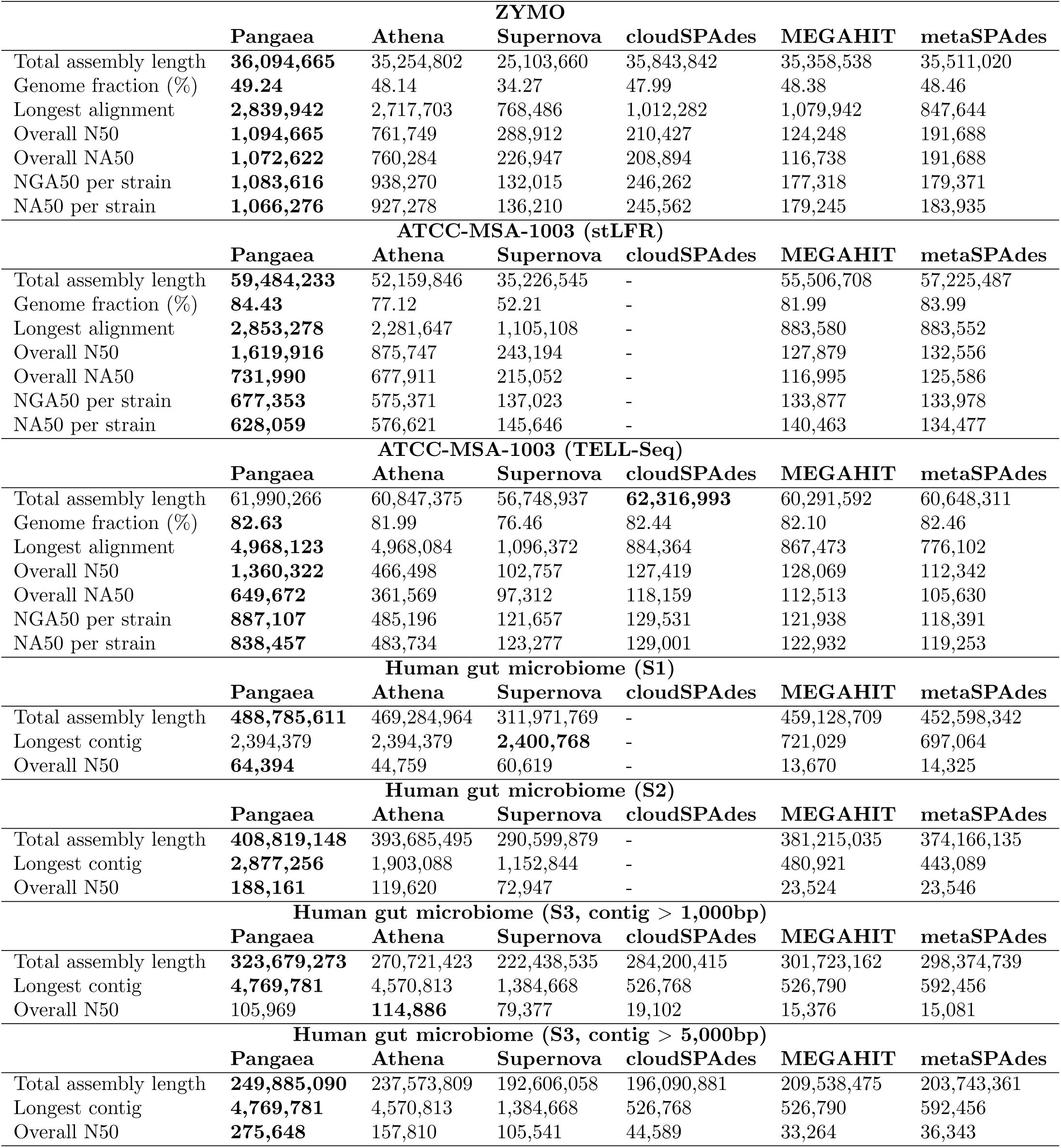
Assembly statistics for different assemblers using the linked-reads datasets on the mock communities and human gut microbiomes. The highest values are in bold.

We simulated 143.04Gb stLFR linked-reads for the ZymoBIOMICS^TM^ Microbial Community Standard II (Log Distribution) [37] (ZYMO; **Supplementary Table 2**; **Methods**), which consists of 10 strains with highly imbalanced abundances (**Supplementary Table 9**). Pangaea also generated substantially higher N50, overall NA50, NA50 per strain, and NGA50 per strain than the other five assemblers on ZYMO (**Table 1**). These observations suggest Pangaea could significantly improve contig continuity using linked-reads with high barcode specificity.

When considering those 15 strains with high abundance (higher than 0.1%) on TELL-Seq linked-reads of ATCC-MSA-1003 (**Supplementary Table 7**), Pangaea still generated significantly higher strain NA50s (**Figure 2 e**) and NGA50s (**Figure 2 h**) than Athena (NA50: p-value = 1.22e-4; NGA50: p-value = 6.10e-5), Supernova (NA50: p-value = 3.05e-4; NGA50: p-value = 3.05e-4), cloudSPAdes (NA50: p-value = 6.10e-5; NGA50: p-value = 6.10e-5), MEGAHIT (NA50: p-value = 6.10e-5; NGA50: p-value = 6.10e-5) and metaSPAdes (NA50: p-value = 6.10e-5; NGA50: p-value = 6.10e-5). A comparable trend was observed on the assemblies of stLFR linked-reads of ATCC-MSA-1003 (**Table 1**; **Figure 2 f and i**). For the 5 strains with the lowest abundance (0.02%), the assemblies of Pangaea had much higher genome fractions than those of Athena (8.12 times on average) and Supernova (54.64 times on average) on stLFR and TELL-Seq linked-reads (**Supplementary Table 8**), indicating more genomic sequences could be assembled by Pangaea for low-abundance microbes.

For ZYMO, Pangaea and Athena always generated better assemblies than the other tools on strains with different abundances (**Supplementary Table 10**). For the two strains (*Escherichia coli* and *Salmonella enterica*) with abundances between 0.01% and 0.1%, Pangaea achieved consistently higher NA50 (1.68 times on average) and NGA50 (1.68 times on average) than Athena (**Supplementary Table 10**). Although all assemblers could not generate contigs covering half of the genome of *Lactobacillus fermentum* (abundance between 0.001% and 0.01%), Pangaea still obtained the highest genome fraction (43.64%), which was substantially higher than the values of the other assemblers considering short-read barcodes (**Supplementary Table 10**). The strains with abundance below 0.001% were unable to be assembled by all assemblers (**Supplementary Table 10**).

### Pangaea generated high-quality assemblies on the human gut microbiomes

We collected DNAs from three healthy Chinese individual stool samples (S1, S2, and S3) and sequenced 136.60Gb (S1), 131.59Gb (S2) and 50.74Gb (S3) stLFR linked-reads for their gut microbiomes (**Supplementary Table 2**; **Supplementary Figure 4**; **Methods**). The assemblies generated by Pangaea achieved the highest total assembly length for all three samples (**Table 1**). Moreover, Pangaea achieved substantially higher N50s than all the other assemblers for both S1 (1.44 times of Athena, 1.06 times of Supernova, 4.71 times of MEGAHIT, 4.50 times of metaSPAdes; **Table 1**) and S2 (1.57 times of Athena, 2.58 times of Supernova, 8.00 times of MEGAHIT, 7.99 times of metaSPAdes; **Table 1**). For S3, Pangaea generated much more sequences than the other assemblers (contig length*>*1Kb; **Methods**), making it unfair to compare their N50 values directly. We transformed their assembly length to be comparable by only considering contigs longer than 5Kb. This led to a significant improvement in the contig N50 of Pangaea, which became the best one (Pangaea: 275.65Kb, Athena: 157.81Kb, Supernova: 105.54Kb, cloudSPAdes: 44.59Kb, MEGAHIT: 33.26Kb, metaSPAdes: 36.34Kb; **Table 1**).

We grouped the contigs into MAGs and identified NCMAGs (**Methods**) to evaluate the performance of metagenome assembly. Pangaea generated NCMAGs (**Figure 3 a**, **e and i**) of 24, 17 and 9 for S1, S2 and S3, which were much more than those generated by Athena (S1: 13, S2: 12, S3: 7), Supernova (S1: 14, S2: 10, S3: 6), cloudSPAdes (only available on S3: 0 due to its low computational efficiency), MEGAHIT (S1: 0, S2: 1, S3: 0) and metaSPAdes (S1: 0, S2: 1, S3: 0). By calculating the number of NCMAGs at different minimum values of N50, we found Pangaea obtained more NCMAGs than the other assemblers at almost all N50 thresholds (**Figure 3 b, f and j**). Pangaea also out-performed the other assemblers with respect to the number of NCMAGs at different maximum read depth thresholds (**Figure 3 c, g and k**). Especially for the NCMAGs with N50s larger than 1Mb, Pangaea achieved substantially more NCMAGs (S1: 8, S2: 4, S3: 5; **Figure 3 d, h and l**) than the other assemblers at all read depth thresholds, while the second best assembler (Athena) only produced 3, 1 and 2 NCMAGs on S1, S2 and S3, respectively (**Figure 3 d, h and l**).

**Figure 3:**
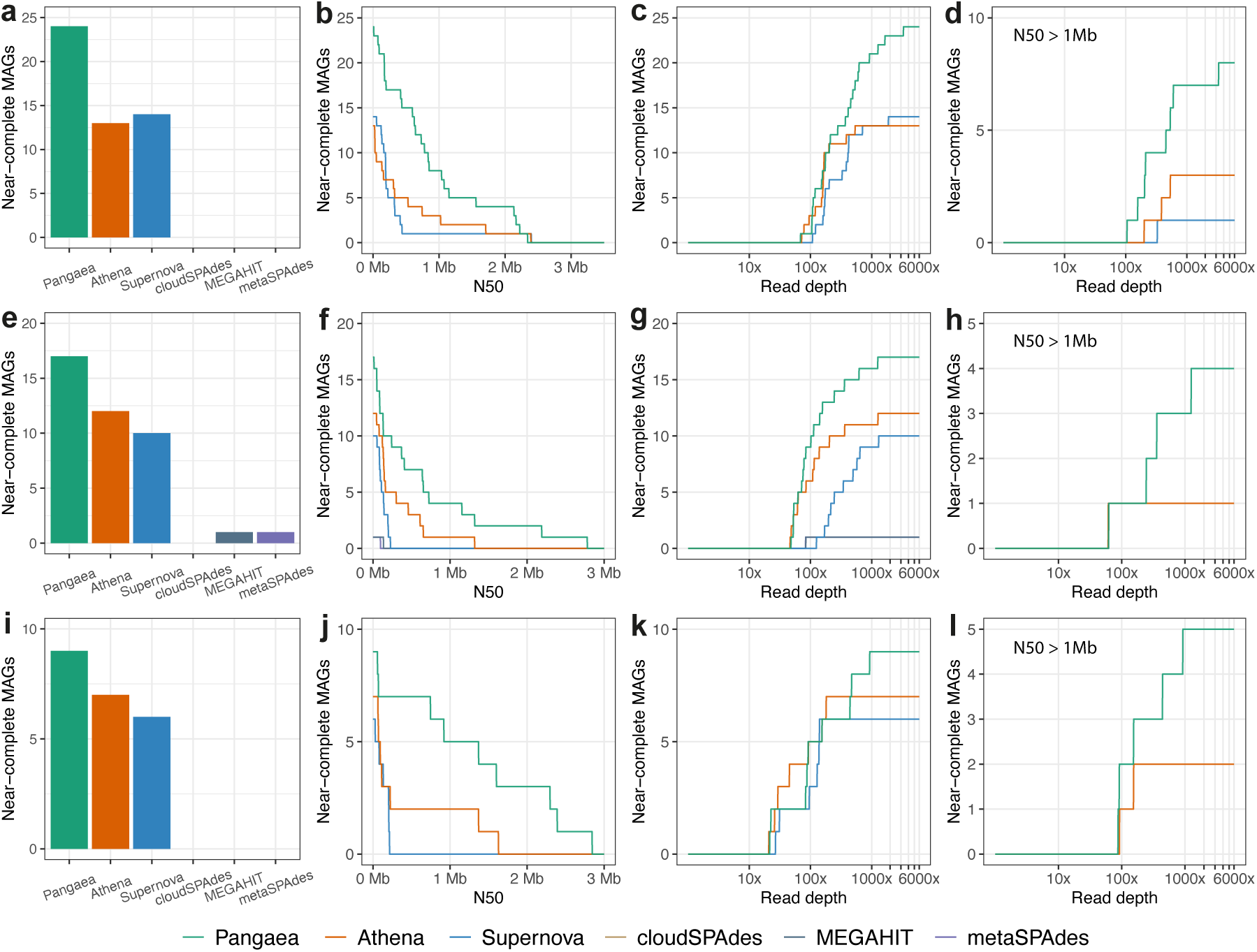
The number of NCMAGs generated by different assembly tools (**a, e, and i**). The number of NCMAGs by thresholding minimum N50 (**b, f, and j**) and maximum read depth (**c-d, g-h, and k-l**). We compared the performances of Pangaea, Athena, Supernova, cloudSPAdes, MEGAHIT and metaSPAdes on stLFR linked-reads from S1 (**a-d**), S2 (**e-h**) and S3 (**i-l**). cloudSPAdes was not available for S1 and S2 because it did not finish assembly within two weeks using 100 threads.

### Pangaea generated high-quality genomes of annotated microbes on linked-reads

We annotated MAGs using Kraken2 [38] with the Nucleotide database of the National Center for Biotechnology Information (**Methods**). Total 61 microbes (S1: 26, S2: 19, S3: 16; **Figure 4**) were annotated from Pangaea’s MAGs; 56 of them (S1: 24, S2: 16, S3: 16; **Figure 4**) achieved the highest N50 and 33 microbes had two-fold higher N50s comparing to the second best assemblers (S1: 16, S2: 8, S3: 9; **Figure 4**). Among the remaining 5 microbes that Pangaea did not achieve the highest N50 (**Figure 4**), Pangaea generated comparable N50s with the best assembler on *Alistipes indistinctus* and *Ruminococcus bicirculans* from S1, and *Bacilli bacterium* from S2; and achieved a lower N50 but substantially higher completeness than Supernova on *Roseburia hominis* (Pangaea: completeness = 96.54%, contamination = 0.48%; Supernova: completeness = 64.91%, contamination = 0.00%; **Supplementary Table 11**) and *uncultured Clostridia* from S2 (Pangaea: completeness = 98.25%, contamination = 7.02%; Athena: completeness = 76.78%, contamination = 1.34%; **Supplementary Table 11**).

**Figure 4:**
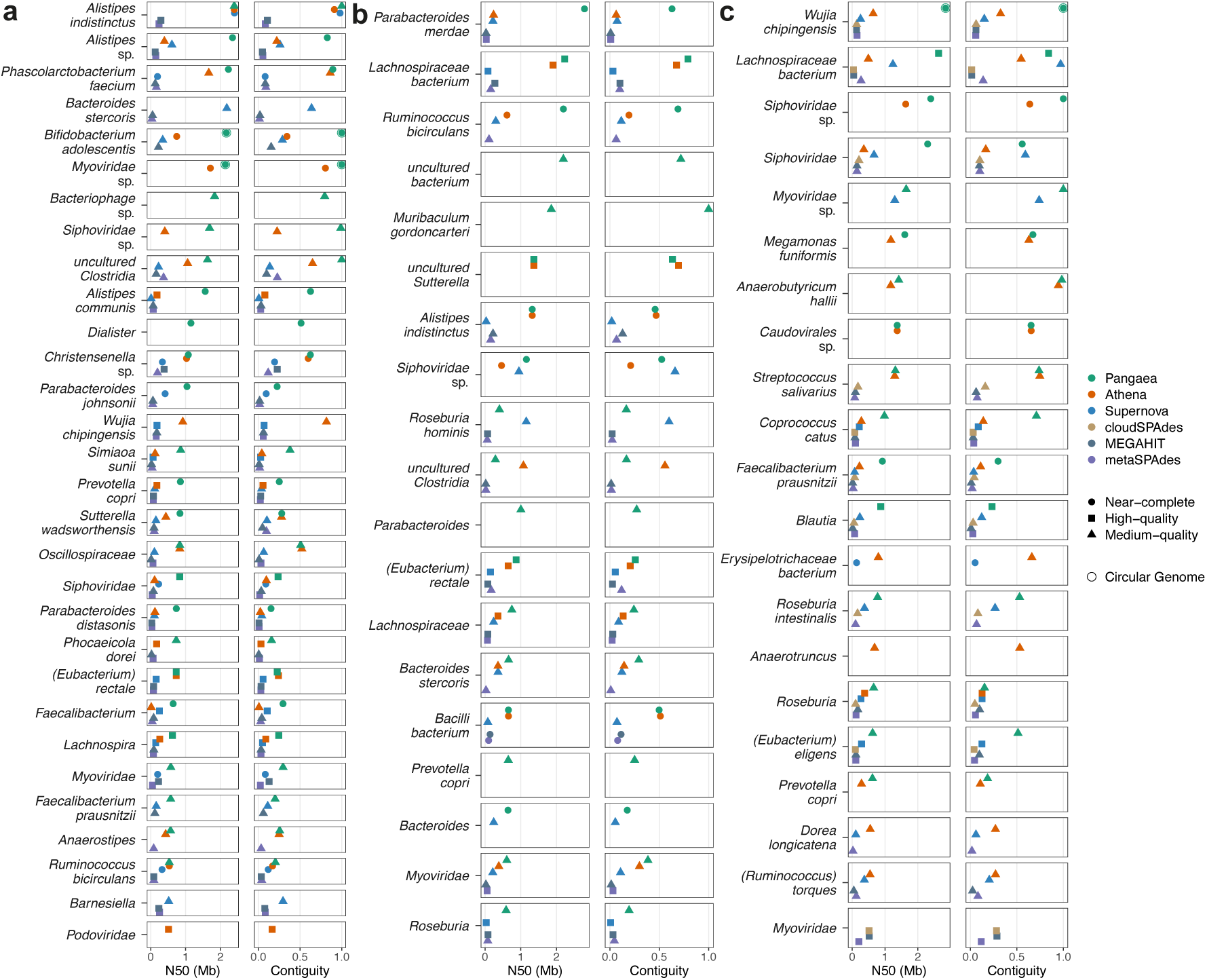
The annotated microbes for the MAGs produced by Pangaea, Athena, Supernova, cloudSPAdes, MEGAHIT and metaSPAdes from S1 (**a**), S2 (**b**) and S3 (**c**). The microbes were shown here if the N50s of their corresponding MAGs were larger than 500Kb by any assembler. If the same microbe was annotated by more than one MAG from the same assembler, the one with the highest N50 was selected. cloudSPAdes was not available for S1 and S2 because it did not finish assembly within two weeks using 100 threads.

Moreover, Pangaea generated more NCMAGs for the annotated microbes. There were 13 microbes (S1: 7, S2: 2, S3: 4; **Figure 4**) that could be assembled as NCMAGs from Pangaea, whereas all the other assemblers only generated MAGs with lower quality or could not generate the matching MAGs. For the three human gut microbiomes, Pangaea identified 17 annotated microbes from the NCMAGs with N50s greater than 1Mb, where Athena, Supernova, cloudSPAdes, MEGAHIT and metaSPAdes only identified 6, 1, 0, 0, and 0 microbes, respectively (**Figure 4**). In addition, Pangaea recognized 6 unique microbes (*Bacteriophage* sp. and *Dialister* from S1, and *uncultured bacterium*, *Muribaculum gordoncarteri*, *Parabacteroides* and *Prevotella copri* from S2) that were not found by any other assemblers, and *Dialister* from S1 was represented by NCMAG from Pangaea (**Figure 4**).

### Strong collinearities between NCMAGs and their closest reference genomes

We compared the NCMAGs that can be annotated as species with their closest reference genomes to evaluate their collinearities (**Methods**). The NCMAGs generated by different assemblers and their closest reference genomes had comparable average alignment identities (Pangaea: 98.16%, Athena: 98.12%, Supernova: 98.31%, MEGAHIT: 98.7%, metaSPAdes: 98.8%) and average alignment frac-tions (Pangaea: 87.9%, Athena: 88.6%, Supernova: 88.4%, MEGAHIT: 88%, metaSPAdes: 90%; **Supplementary Table 12**), while Pangaea produced significantly more species-level NCMAGs than the other assemblers (Pangaea: 29, Athena: 21, Supernova: 14, cloudSPAdes: 0, MEGAHIT: 1, metaSPAdes: 1; **Supplementary Table 12**).

The NCMAGs assembled by Pangaea had high collinearities with their closest reference genomes (**Figure 5**; **Supplementary Figure 5**). Some NCMAGs showed inversions and rearrangements in comparison to the reference sequences, including *Alistipes communis* (S1; **Supplementary Figure 5 a**), *Desulfovibrio desulfuricans* (S1; **Supplementary Figure 5 c**) and *Siphoviridae* sp. (S2; **Figure 5 g**). Pangaea assembled NCMAGs for *Siphoviridae* sp. from both S2 and S3 (**Figure 5 g and h**). The two NCMAGs had comparable total sequence lengths (S2: 2.20Mb and S3: 2.39Mb; **Supplementary Table 11**) and better N50 was achieved in S3 (N50: 1.16Mb for S2 and 2.39Mb for S3; **Supplementary Table 11**). Note that the NCMAG of *Siphoviridae* sp. from S3 is a single-contig NCMAG, but it’s not a circular contig which might be because it does not include all single-copy genes (completeness: 91.28%; **Supplementary Table 11**).

Pangaea could generate NCMAGs for annotated species with better quality and larger N50 values than the other assemblers, such as *Alistipes* sp. from S1 and *Collinsella aerofaciens* from S2 (**Figure 5 d and e**; **Supplementary Table 11**). The evaluation of the read depths and GC-skew of the MAGs revealed that Pangaea recovered the regions with extremely low read depths and high GC-skew, such as the region at approximately 480Kb of *Faecalibacterium prausnitzii* from S3 (**Supplementary Figure 5 q**). This indicates Pangaea has the potential to reconstruct hard-to-assemble genomic regions.

### Pangaea generated complete and circular MAGs using linked-reads

We examined if there existed complete and circular genomes in NCMAGs based on the circularization module in Lathe [39] (**Methods**). We found that only Pangaea generated three circular NCMAGs, which were annotated as *Bifidobacterium adolescentis* (S1), *Myoviridae* sp. (S1) and *Wujia chipingensis* (S3). For each of the three microbes, Pangaea produced a gapless contig with perfect collinearity with the closest reference genomes (**Figure 5 a, b and i**).

Athena generated three and two contigs for *B. adolescentis* (from S1) and *Myoviridae* sp. (from S1), with substantially lower contig N50 values than the contigs obtained by Pangaea for these two species (*B. adolescentis*: Pangaea = 2,167.94Kb, Athena = 744.54Kb; *Myoviridae* sp.: Pangaea = 2,137.66Kb, Athena = 1,709.63Kb; **Supplementary Table 11**). Supernova, MEGAHIT and metaSPAdes could only generate incomplete MAGs or could not assemble these two species, and the completeness of their candidate MAGs was significantly lower than that of MAGs generated by Pangaea (**Supplementary Table 11**). For *W. chipingensis* from S3, Pangaea was the only assembler that obtained NCMAG and got a significantly higher contig N50 than the other assemblers (Pangaea: 2,846.41Kb, Athena: 639.65Kb, Supernova: 254.31Kb, cloudSPAdes: 135.56Kb, MEGAHIT: 134.14Kb, metaSPAdes: 146.44Kb; **Supplementary Table 11**).

### Pangaea generated high-quality assemblies on short-reads with virtual barcodes from long-reads

We evaluated the generalizability of Pangaea on hybrid assembly in terms of attaching virtual barcodes to short-reads. We attached identical barcodes to short-reads if they were aligned to the same long-reads, which is another way to offer long-range connectivity of short-reads (**Methods**). We used the short-reads alignment with 1.27Gb PacBio CLR [13] (ATCC-MSA-1003) and 2.48Gb Nanopore GridION (ZYMO) [37] long-reads (**Supplementary Table 2**) to substitute the barcodes of TELL-Seq (ATCC-MSA-1003) and stLFR (ZYMO) linked-reads. We adjusted the workflow of Pangaea to integrate with hybridSPAdes (namely Pangaea-Hybridspades) and OPERA-MS (namely Pangaea-Operams; **Methods**). We compared Pangaea-Hybridspades and Pangaea-Operams to hybridSPAdes, OPERA-MS, Athena and metaSPAdes, and observed that virtual barcodes could prominently increase contig N50 and NA50 (**Table 2**; **Figure 6 a**). For example, the N50 and NA50 of Pangaea-Hybridspades were 1.7 and 1.31 times higher than hybridSPAdes while the corresponding values of OPERA-MS were increased by 4.43 and 3.58 folds (**Table 2**).

**Figure 5:**
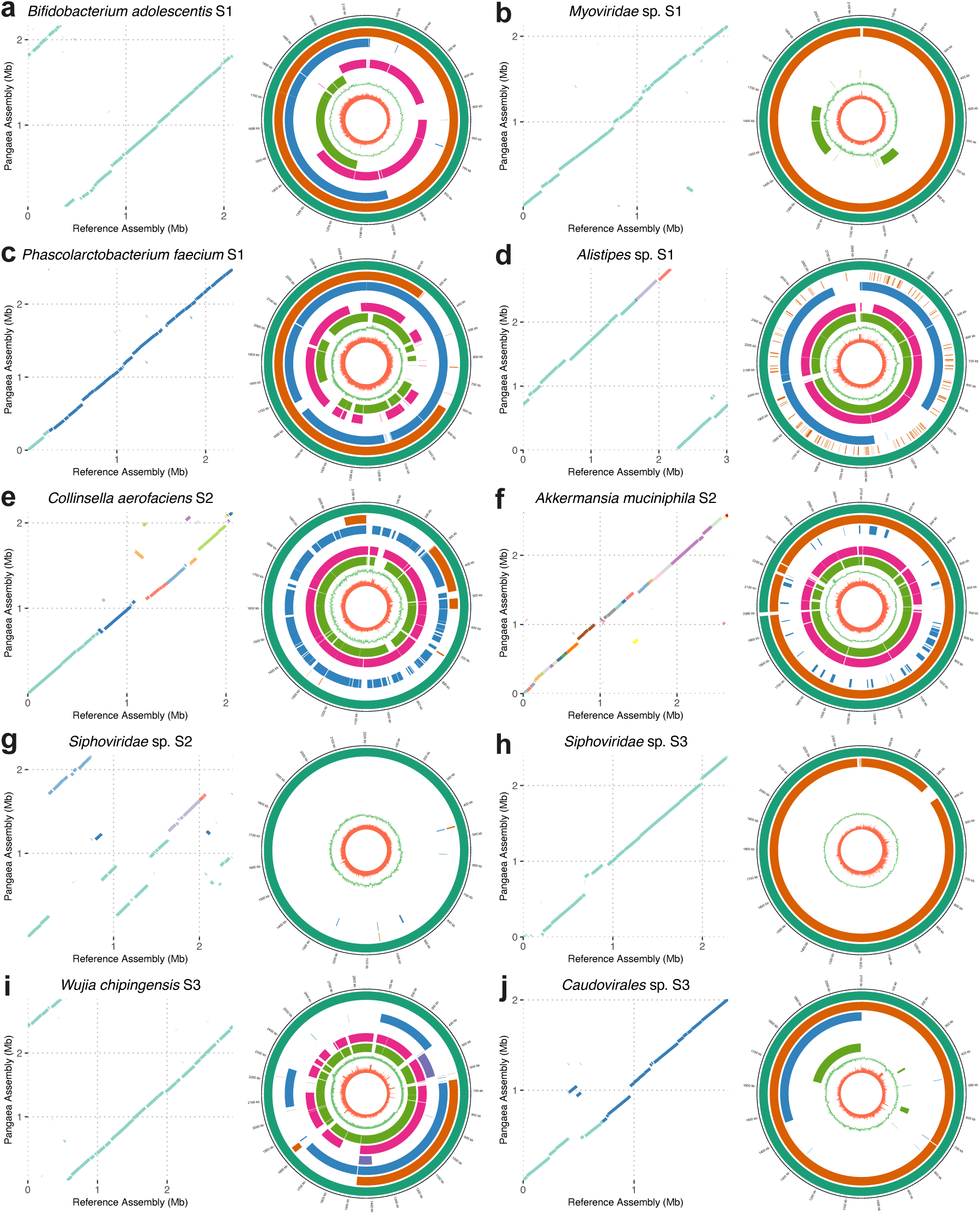
Genome collinearity analysis between the selected NCMAGs produced by Pangaea and their closest reference genomes (dot plots), and comparison of the corresponding MAGs produced by different assemblers (circos plots) from S1, S2 and S3. **a, b and i** are species for which Pangaea obtained complete and circular MAGs. Colors are used in the dot plots to distinguish different contigs. The eight rings in the circos plots from outside to inside denote the Pangaea MAGs (dark green), Athena MAGs (orange), Supernova MAGs (blue), cloudSPAdes MAGs (purple), MEGAHIT MAGs (pink), metaSPAdes MAGs (light green), GC-skew of Pangaea MAGs, and read depth of Pangaea MAGs, respectively. If the same species was annotated by more than one MAG from the same assembler, the one with the highest N50 is shown here. The remaining NCMAGs produced by Pangaea are shown in **Supplementary Figure 5**.

**Figure 6:**
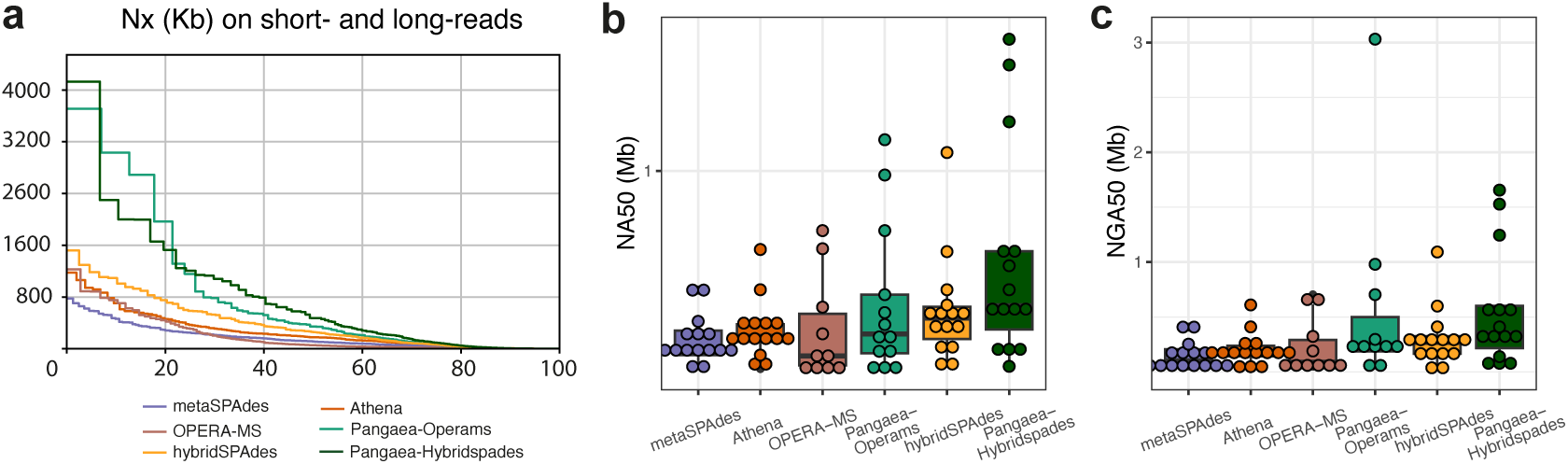
Nx with x ranging from 0 to 100 (**a**), NA50 of the 15 strains with abundance *>* 0.1% (**b**), and NGA50 of the 15 strains with abundance *>* 0.1% (**c**), assembled by different assemblers using short-reads (metaSPAdes), short-reads with virtual barcodes (Athena, Pangaea-Operams and Pangaea-Hybridspades), and using short- and long-reads (OPERA-MS and hybridSPAdes) on ATCC-MSA-1003 mock community.

**Table 2:**
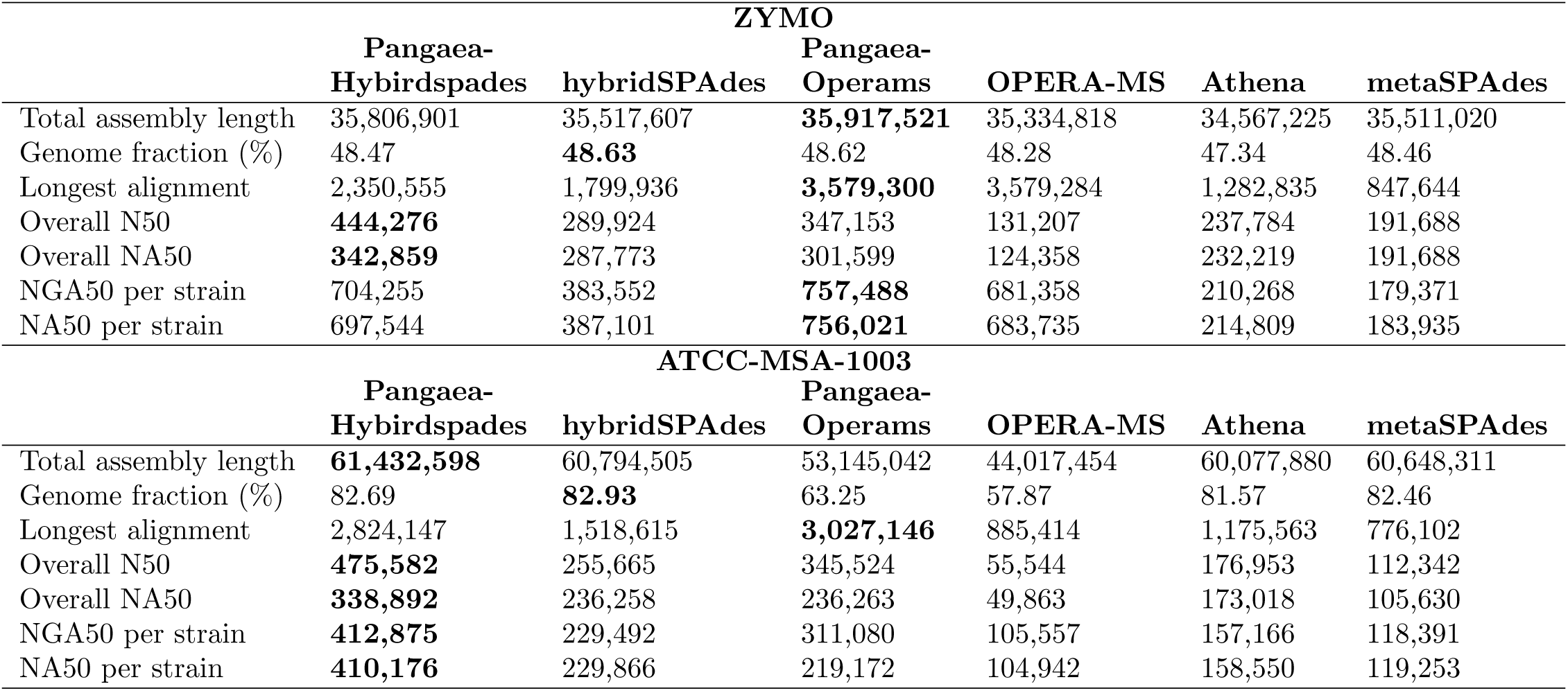
Assembly statistics for different assemblers using short-reads with virtual barcodes from long-reads on the ZYMO and ATCC-MSA-1003. The highest values are in bold.

For ATCC-MSA-1003, Pangaea-Hybridspades achieved the highest N50s (average N50: 903.64Kb) and NA50s (average NA50: 701.48Kb; **Supplementary Table 13**) for the 10 high- and medium-abundance strains with abundance higher than 1%. It also generated significantly higher NA50s and NGA50s than the other assemblers for the 15 strains with abundances higher than 0.1% (**Figure 6 b and c**). For the 5 strains with the lowest abundance (0.02%), all assemblers got poor NA50s and NGA50s; Pangaea-Operams achieved much more sequences from the contigs longer than 10Kb compared to OPERA-MS (Pangaea-Operams: 141.76Kb, OPERA-MS: 64.19Kb; **Supplementary Table 13**), suggesting Pangaea could improve contig continuity for low-abundance microbes.

For ZYMO, Pangaea-Hybridspades also generated significantly higher average NA50 (Pangaea-Hybridspades: 1.98Mb, hybridSPAdes: 887.00Kb; **Supplementary Table 13**) and NGA50 (Pangaea-Hybridspades: 1.98Mb, hybridSPAdes: 887.00Kb; **Supplementary Table 13**) for the two strains with abundances higher than 1% (*Listeria monocytogenes* and *Pseudomonas aeruginosa*) than hybridSPAdes. For the two strains (*Escherichia coli* and *Salmonella enterica*) with abundance between 0.01% and 0.1%, Pangaea-Hybridspades and hybridSPAdes had comparable contig NA50 and NGA50; while Pangaea-Operams produced better NA50 (1.66 times on average) and NGA50 (1.71 times on average) than OPERA-MS (**Supplementary Table 13**). All the assemblers generated low-quality assemblies for the strain with an abundance between 0.001% and 0.01% (*Lactobacillus fermentum*). Pangaea-Hybridspades (19.77Kb) and Pangaea-Operams (54.91Kb) achieved much more sequences from the contigs longer than 10Kb than hybridSPAdes (13.69Kb) and OPERA-MS (37.41Kb), respectively (**Supplementary Table 13**). The strains with abundances below 0.001% cannot be assembled for all assemblers (**Supplementary Table 13**). These observations demonstrate that Pangaea could improve the assemblies for both high-abundance and low-abundance microbes by leveraging virtual barcodes from long-reads.

### Evaluation of running time and maximum memory usage

We compared the computational performance (CPU time, Real time, and Maximum Resident Set Size [RSS]) of the benchmarked assemblers on ZYMO (**Figure 7**; **Methods**). Both Athena and Pangaea required the assemblies from the other tools, so we only considered their additional processing time and memory. MEGAHIT was the fastest assembler with the lowest maximum RSS (**Figure 7**). metaSPAdes and cloudSPAdes consumed substantially higher CPU times than the other assemblers (**Figure 7 a**). The real time (wall clock time) used by metaSPAdes, cloudSPAdes, Supernova, Athena and hybridSPAdes were comparable and significantly higher than those consumed by MEGAHIT, Pangaea, OPERA-MS, Pangaea-Operams and Pangaea-Hybridspades (**Figure 7 b**). Supernova used the highest maximum RSS on ZYMO (**Figure 7 c**). The maximum RSSs used by metaSPAdes, cloudSPAdes, and hybridSPAdes were close to each other, and much higher than those needed by the other assemblers (**Figure 7 c**). These results revealed Pangaea could improve the assembly quality of the existing assemblers in a reasonable time and using a relatively low maximum RSS.

**Figure 7:**
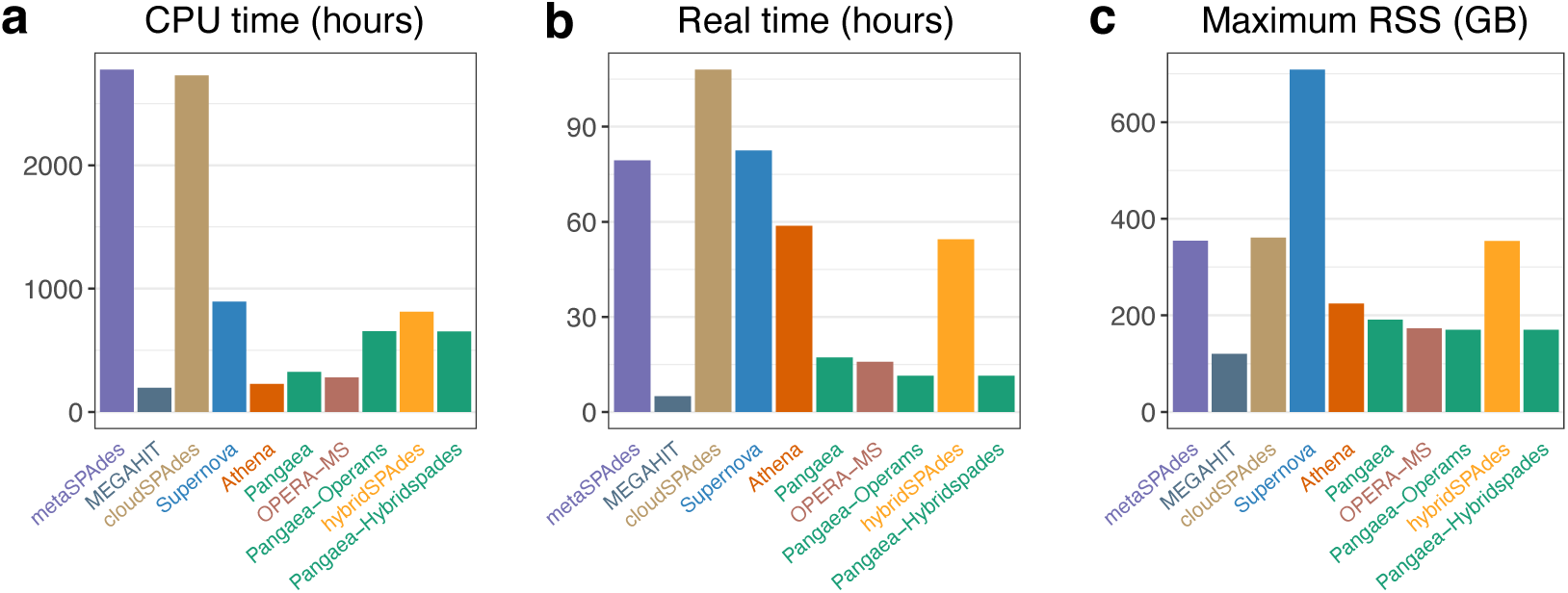
The CPU time (**a**), real time (**b**) and maximum resident set size (RSS; **c**) of the assemblers using short-reads (metaSPAdes and MEGAHIT), linked-reads (cloudSPAdes, Supernova, Athena and Pangaea) and using short- and long-reads (OPERA-MS, Pangaea-Operams, hybridSPAdes and Pangaea-Hybridspades) on ZYMO mock community.

## Discussion

Short-read sequencing has proven to be an important approach for analyzing human gut microbiota from large sequencing cohorts. However, its lack of long-range DNA connectivity makes assembling conserved sequences, intra- and inter-species repeats, and ribosomal RNAs (rRNAs) difficult [5]. It has limitations in producing complete microbial genomes and long-read sequencing is relatively expensive to be applied to large cohorts. Cost-effective linked-read sequencing technologies, which attach barcodes to short-reads to provide long-range DNA connectivity, have achieved great success in improving contig continuity in metagenome assembly [14, 19]. Unlike 10x linked-reads, stLFR [17] and TELL-Seq linked-reads [18] have high barcode specificity, but a dedicated assembler that could make full use of this characteristic to improve metagenome assembly is lacking. Besides linked-reads, the long-range connectivity of short-reads could also be provided by virtual barcodes from other independent sequencing technologies (e.g., long-reads).

In this study, we developed Pangaea to improve metagenome assembly by leveraging long-range connectivity from linked-reads and long-reads. It considers the co-barcoded reads as long DNA fragments and extracts their *k*-mer histograms and TNFs for co-barcoded read binning. This strategy significantly reduces the complexity of metagenomic sequencing data and makes the assembly more efficient. Because sequence clustering is sensitive to data sparsity and noise, Pangaea represents the input features in a low-dimensional latent space using VAE. We also designed a weighted sampling strategy to generate a balanced training set for microbes with different abundances. This module primarily advantages microbes with high- and medium-abundance, because they are more robust to mis-binning and usually have sufficient data for assembly in the corresponding bins. Pangaea adopts a multi-thresholding reassembly strategy to rescue the reads from low-abundance microbes. It eliminates short-reads from high-abundance microbes in the assembly graph gradually to differentiate the spurious edges from low-abundance microbes and sequencing errors. In the third module, we merged the assemblies from different strategies due to their complementary nature of each other.

The previous study [40] showed that co-assembly with multiple samples could improve completeness and decrease the contamination of MAGs. However, the co-assembly of metagenomes derived from large samples poses a practical challenge due to time and computational limitations. Because the sequencing data from all the individual samples need to be merged before co-assembly. Read binning is a strategy that enables co-assembly on large datasets by producing smaller and simpler subsets of reads, thereby facilitating the assembly process. Several studies have attempted to apply read binning to short-read metagenomic sequencing [29, 28, 27], but it is exceedingly difficult in practice. The fragments of short-reads are too short to allow the extraction of stable sequence abundance and composition features from the individual reads. Existing read binning tools have to identify the overlap between each pair of reads for binning. However, the millions or even billions of short-reads make the overlap-based read binning algorithm extremely slow and highly memory intensive. Overlap Graph-based Read clustEring (OGRE) [27] was developed to improve the computational performance of read binning, but it still consumed 2,263 CPU hours even for the low-complexity dataset of CAMI [27]. We evaluated OGRE on stLFR linked-reads of ATCC-MSA-1003 (664.77M read pairs) and observed that OGRE crashed due to insufficient memory if 100 threads were applied. If fewer threads were applied, the binning time would become extremely long (more than two weeks). Pangaea with 100 threads only took 64.06 hours in real time, 514.63 hours in CPU time, and consumed 281.99GB of maximum RSS to group and assemble this linked-reads dataset.

Long-read sequencing has received increasing attention due to its ability to generate complete microbial genomes from complex communities. However, it is limited by a relatively high cost for large cohort studies. In contrast, linked-read and hybrid sequencing (deep short-read and shallow long-read sequencing) techniques are cost-effective. Linked-read sequencing only requires a tiny amount of input DNA, and can thus be a complementary solution to long-read sequencing. In our experiments, we found that long-read assemblies had 60.98% fewer NCMAGs than linked-read assemblies from Pangaea (**Supplementary Note 4**), indicating that some microbes might be lost due to insufficient long-read sequencing depth. Similar observations have been reported in previous studies [39]. Although stLFR and TELL-Seq linked-reads had high barcode specificity in ATCC-MSA-1003, we observed that a considerable fraction of barcodes still contained more than one fragment (stLFR = 37.02%, TELL-Seq = 72.95%), which could complicate the deconvolution of barcodes for the existing linked-read assemblers. We believe that further protocol improvement for these technologies (e.g. increasing the number of beads) may further improve their metagenome assembly performance.

## Methods

### DNA preparation and sequencing for linked-read sequencing

For ATCC-MSA-1003, the microbial DNAs were extracted directly from the 20 Strain Staggered Mix Genomic Material (ATCC MSA-1003) without size selection using a QIAamp DNA stool mini kit (Qiagen, Valencia, CA, USA). For the human gut microbiomes, microbial DNAs from stool samples of three individuals (S1, S2, and S3) were extracted using the QIAamp DNA stool mini kit (Qiagen) and size-selected using a BluePippin instrument targeting the size range of 10-50 Kb according to the manufacturer’s protocol. The stLFR libraries were prepared using the stLFR library prep kit (16 RXN), followed by 2*×*100 paired-end short-read sequencing using BGISEQ-500. The TELL-Seq library for ATCC-MSA-1003 was prepared using the TELL-Seq Whole Genome Sequencing library prep kit, followed by 2×146 paired-end sequencing on an Illumina sequencing system.

### Simulate stLFR linked-reads for ZYMO

We downloaded reference genomes of the 10 strains of the ZymoBIOMICS^TM^ Microbial Community Standard II (Log Distribution) [37], and used LRTK (v1.7) [41] to simulate stLFR linked-reads with the same strain composition as the ZYMO mock community.

### Generate virtual barcodes to short-reads from long-reads

We attached the same long-read indexes as virtual barcodes to short-reads if they were aligned to the same long-read. We mapped the short-reads to long-reads using BWA-MEM (v0.7.17) [42] and removed the spurious alignments if the minimum aligned nucleotides were below 60bps. If a short-read was aligned to multiple long-reads, we randomly chose one of the long-read indexes as its barcode.

### Extract *k* -mer histogram and TNFs from co-barcoded reads

We extracted *k*-mer histograms and TNFs from the co-barcoded reads if their total lengths were longer than 2Kb to ensure feature stability. A *k*-mer histogram was calculated based on global *k*-mer occurrences and could reflect the abundance features of the microbial genome [33]. We adopted *k* = 15 as used in the previous studies [33, 43], and calculated the global 15-mer frequencies using the whole dataset. We removed 15-mers with frequencies higher than 4,000 (to avoid repetitive sequences) and divided the global frequency distribution into 400 bins with equal sizes (the *i^th^* bin denoted frequencies between 10 *∗ i −* 10 and 10 *∗ i*). We collected the co-barcoded reads for each barcode and divided these reads into 15-mers, which were assigned to the 400 bins based on their global frequencies. We calculated the number of 15-mers allocated to each bin and generated a count vector with 400 dimensions as the *k*-mer histogram of the specific barcode. A TNF vector was constructed by calculating the frequencies of all 136 non-redundant 4-mers from co-barcoded reads. The *k*-mer histogram and TNF vector were normalized to eliminate the bias introduced by the different lengths of co-barcoded reads.

### Binning co-barcoded reads with a VAE

The normalized *k*-mer histogram (*X_A_*) and TNF vector (*X_T_*) were concatenated into a vector with 536 dimensions as the input to a VAE (**Figure 1 b**; **Supplementary Note 2**). The encoder of VAE consisted of two fully connected layers with 512 hidden neurons, and each layer was followed by batch normalization [44] and a dropout layer (P = 0.2) [45]. The output of the last layer was fed to two parallel latent layers with 32 hidden neurons for each to generate *µ* and *σ* for a Gaussian distribution 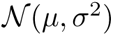, from which the embedding *Z* was sampled. The decoder also contained two fully connected hidden layers of the same size as the encoder layers to reconstruct the input vectors 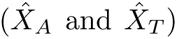 from the latent embedding Z. We applied the *softmax* activation function on 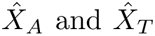 to achieve the normalized output vectors, because the input features *X_A_* and *X_T_* were both normalized. The loss function (*Loss*) was defined as the weighted sum of three components: the reconstruction loss of *k*-mer histogram (*L_A_*), the reconstruction loss of TNF vectors (*L_T_*), and the Kullback-Leibler divergence loss (*L_KL_*) between the latent and the prior standard Gaussian distributions. We adopted cross-entropy loss for *L_A_* and *L_T_* to deal with probability distributions, and formularized the loss terms as follows:

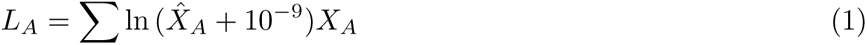

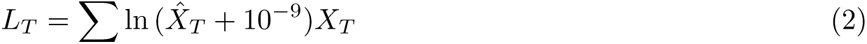

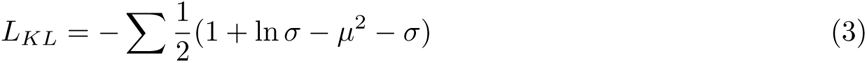

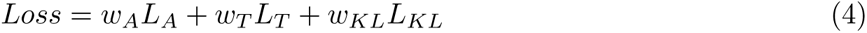

where the weights of the three loss components were *w_A_* = *α/* ln(*dim*(*X_A_*)), *w_T_* = (1*−α*)*/* ln(*dim*(*X_T_*)), and *w_KL_* = *β/dim*(*Z*). We adopted 0.1 and 0.015 for *α* and *β*, respectively (**Supplementary Note 3**). The VAE was trained with early stopping to reduce the training time and avoid overfitting. We used the RPH-kmeans [34] algorithm with random projection hashing to group the co-barcoded reads using their latent embeddings obtained from *µ*. We set the number of clusters equal to or smaller than the actual number of microbes in the microbial community. For the two mock communities of ZYMO and ATCC-MSA-1003, the numbers of clusters were set to 10 and 15, respectively. For all three real human gut microbiomes, we used 30 as the number of clusters.

### Weighted sampling for VAE training

We designed weighted sampling to balance the training set of co-barcoded reads from microbes with different abundances. Considering that the highest value of *X_A_* is negatively related to the abundance of the co-barcoded reads (**Supplementary Figure 1**), we used a heuristic function *max*(*X_A_*)^2^ as the sampling weight for the barcode of the co-barcoded reads. The square was to make the low-abundance microbes have a much higher sampling weight. The calculated sampling weights were automatically used by the WeightedRandomSampler of PyTorch to construct a balanced training dataset.

### Multi-thresholding reassembly for low-abundance microbes

For co-barcoded read binning, we assembled reads in each cluster independently using MEGAHIT (v1.2.9) [31]. We designed a multi-thresholding reassembly strategy (**Figure 1 b**) to improve the assembly qualities of low-abundance microbes by recollecting the reads from the low-abundance microbes that were misclustered into different bins using read depth thresholds. To calculate the read depth of the contigs assembled from each read cluster (denoted as contigs*_bin_*), we aligned the input reads to contigs*_bin_* using BWA-MEM (v0.7.17) [42] and calculate the read depth for each contig using the “jgi summarize bam contig depths” program in MetaBat2 (v2.12.1) [46]. To collect the reads of low-abundance microbes, we extracted the reads that could not be mapped to the high-depth contigs (with read depth *> t_i_*) in contigs*_bin_* and assembled them using the standard short-read assembler, metaSPAdes (v3.15.3) [30]. This step can be substituted by MEGAHIT to reduce the running time, and can also integrate available contigs to guide the path resolution in metaSPAdes to make the assembly of low-abundance microbes more efficient (e.g., using the contigs assembled by metaSPAdes from the whole read dataset as the input to “--untrusted-contigs” of metaSPAdes, which is used in our experiments and optional for Pangaea). We repeated this procedure with a range of thresholds (*T* = {*t_i_|i* = 1, 2*, ..*.}) producing contigs*_low_*. We chose *T* = {10, 30} for all the experiments, which worked well for both the mock communities and the human gut microbiomes.

### Ensemble assembly

We use ensemble assembly to avoid incomplete metagenome assembly caused by the mis-binning of previous modules. For linked-read assembly, the ensemble strategy includes two steps: (i) we use contigs*_bin_* (contigs assembled from each read bin), contigs*_low_* (contigs from multi-thresholding reassembly), contigs*_local_* (contigs from the local assembly of Athena) and contigs*_ori_* (contigs assembled from short-reads by metaSPAdes [v3.15.3]) using an OLC-based assembler, metaFlye (v2.8) with the “--subassemblies” parameter; (ii) quickmerge (v0.3) [47] was used to merge the contigs from step (i) and Athena contigs. For the assembly of short-reads with virtual barcodes, we substituted the metaSPAdes in step (i) and Athena in step (ii) with the corresponding hybrid assemblies (contigs generated from hybridSPAdes [v3.15.3] [16] or OPERA-MS [v0.8.3] [15]). Step (ii) is optional for linked-read assembly since Athena is already integrated in Step (i).

### Detecting circular contigs

We adopted the circularization module of Lathe [39] to detect circular contigs in all the assemblies. The module needs long-reads as input which is not available for linked-read assembly, so we modified the alignment and assembly parameters in the circularization module to accept contigs as input, and merged the contigs*_ori_*, contigs*_bin_*, contigs*_low_*, and contigs*_local_* as “pseudo long-reads” for running it.

### Reconstructing physical long fragments based on reference genomes

We reconstructed the physical long fragments from linked-reads of ATCC-MSA-1003 to calculate *N_F/B_*. The linked-reads were mapped to the reference genomes using BWA-MEM (v0.7.17) [42] with option “-C” to retain the barcode information in the alignment file, followed by sorting based on read alignment coordinates using SAMtools (v1.9) [48]. We connected the co-barcoded reads into long fragments if their coordinates were within 10Kb on the reference genome. Each fragment was required to include at least two read pairs and to be no shorter than 1Kb.

### Metagenome assembly of the other assemblers on different datasets

The 10x, stLFR, and TELL-Seq sequencing datasets were demultiplexed to generate raw linked-reads using Long Ranger (v2.2.0) [49], stLFR read demux (Git version 3ecaa6b) [17] and LRTK (Git version 28012df) [41], respectively. The linked-reads were assembled using metaSPAdes (v3.15.3) [30], MEGAHIT (v1.2.9) [31], cloudSPAdes (v3.12.0-dev) [30], Athena (v1.3) [14] and Supernova (v2.1.1) [20]. Supernova does not accept raw stLFR linked-reads as input, so we applied stlfr2supernova pipeline (https://github.com/BGI-Qingdao/stlfr2supernova pipeline; Git version 95f0848) to convert the barcode format of stLFR. The scaffolds produced by Supernova were broken into contigs at successive “N”s longer than 10 before evaluation. The datasets with both short- and long-reads were assembled using OPERA-MS (v0.8.3) [15], hybridSPAdes (v3.15.3) [16], metaSPAdes (v3.15.3, only short-reads) and Athena (v1.3, short-reads with virtual barcodes). The PacBio CLR long-reads from S1 and S2 were assembled using metaFlye (v2.8) [50] with the “--pacbio-raw” parameter. All the assemblers were run with default parameters. For measuring computational performance, we set the threads of all the assemblers to 100 (if the assembler had this option), and used the command “/usr/bin/time -v” to report the system time, user time, and maximum RSS consumed by the programs.

### Benchmarking on the mock microbial communities

The reference genomes of ATCC-MSA-1003 and ZYMO were downloaded from the NCBI reference databases (**Supplementary Table 1**) and the previous study on ZYMO [37], respectively. The contigs assembled for the two mock communities were assessed using MetaQUAST (v5.0.2) [51], with the option “-m 1000 --fragmented --min-alignment 500 --unique-mapping” to enable the alignment of fragmented reference genomes and discard ambiguous alignments. The p-values of differences in the NA50 and NGA50 of different assemblers were obtained using the Wilcoxon signed-rank test performed by the *wilcox.test* function of R with “paired = TRUE”.

### Contig binning and MAG quality evaluation

We aligned the linked-reads (or short-reads) to the contigs using BWA-MEM (v0.7.17) [42] and calculated the read depths using “jgi summarize bam contig depths” in MetaBat2 (v2.12.1) [46]. The contigs with read depths were binned into MAGs using MetaBat2 (v2.12.1) with default parameters. CheckM (v1.1.2) [52] was used to report the completeness and contamination of the MAGs. ARAGORN (v1.2.38) [53] and barrnap (v0.9) [54] were used to annotate the transfer RNAs (tRNAs) and rRNAs (5S, 16S, and 23S rRNAs), respectively. According to standard criteria of the minimum information about MAGs [55], we classified the MAGs into near-complete (completeness *>* 90%, contamination *<* 5%, and could be detected 5S, 16S, and 23S rRNAs, and at least 18 tRNAs), high-quality (completeness *>* 90%, and contamination *<* 5%), medium-quality (completeness *≥* 50%, and contam- ination *<* 10%), and low-quality (the other MAGs).

### Annotation of the MAGs and the closest reference genomes

The contigs were annotated using Kraken2 (v2.1.2) with the custom database built from the NT database of NCBI (Aug 20, 2022). We used the “--fast-build” option of kraken2-build to reduce the database construction time. Subsequently, the “assign species.py” script from “metagenomics workflows” [14, 39] was used to annotate MAGs as species (if the fraction of contigs belonging to the species was more than 60%) or genus (otherwise) based on contig annotations. The closest reference genomes of the NCMAGs that can be annotated at species-level were identified using GTDB-Tk (v2.1.0; database version r207) [56], which also reported the alignment identities and alignment fractions between them.

## Acknowledgements

We thank Tom Chen and Yong Wang from Universal Sequencing Technology for providing the TELL-Seq sequencing data of the ATCC-MSA-1003 mock community. We thank Arend Sidow for his comments to improve the manuscript’s language and structure. We also thank the Research Committee of Hong Kong Baptist University and the Interdisciplinary Research Clusters Matching Scheme for their kind support of this project.

## Abbreviations

MAG: Metagenome-Assembled Genome
NCMAG: Near-Complete Metagenome-Assembled Genome
OGRE: Overlap Graph-based Read clustEring
PacBio CLR: PacBio Continuous Long-Reads
RSS: Resident Set Size
rRNA: Ribosomal RNA
stLFR: single-tube Long Fragment Read
TELL-Seq: Transposase Enzyme-Linked Long-read Sequencing
TNF: TetraNucleotide Frequency
tRNA: Transfer RNA
UST: Universal Sequencing Technology
VAE: Variational AutoEncoder

## Availability of data and materials

The 10x linked-reads of the ATCC-MSA-1003 mock community were downloaded from NCBI SRA accession SRR12283286. The stLFR and TELL-Seq sequencing data of ATCC-MSA-1003 was uploaded to NCBI BioProject PRJNA875547. The stLFR sequencing data of the three human gut microbiomes was deposited in China National GeneBank (CNGB) project CNP0003432. The PacBio CLR long-reads of ATCC-MSA-1003, S1 and S2 were downloaded from NCBI SRA accessions SRR12371719, SRR19505636 and SRR19505632, respectively. The ONT long-reads of ZYMO were available from the NCBI SRA accession ERR3152366. Codes of Pangaea and all the command lines are available at https://github.com/ericcombiolab/Pangaea. The version of the source code used in the manuscript was uploaded to https://doi.org/10.5281/zenodo.7996172.

## Ethics approval and consent to participate

Not applicable

## Consent for publication

Not applicable

## Authors’ contributions

LZ conceived the study. ZMZ, LZ and JX designed the Pangaea algorithms. ZMZ and JX implemented the Pangaea software. LZ and ZMZ conceived the experiments. ZMZ and JX conducted the experiments. ZMZ, LZ and JX analyzed the results. HBW drew and analyzed the circos plots. CY generated and analyzed the statistics of the three types of linked-reads. ZMZ and LZ wrote the manuscript. XDF, YFH, ZY, YC and LJH sequenced the stLFR linked-reads. APL revised the paper and supported the project. All authors reviewed the manuscript.

## Additional Files

**Supplementary Text and Figures**

Supplementary Notes 1-4, Supplementary Figures 1-5, and Supplementary Tables 1-3, 6, and 9.

**Supplementary Table 4**

The impact of VAE-based binning algorithm adopted by Pangaea on the final assembly results using TELL-Seq linked-reads of the ATCC-MSA-1003 mock community.

**Supplementary Table 5**

The impact of different ranges of abundance thresholds (Ts) on the final assembly using the TELL-Seq linked-reads of the ATCC-MSA-1003 mock community.

**Supplementary Table 7**

The per-strain NA50 and NGA50 generated by different assembly tools for the 15 strains with abundances higher than 0.1% using TELL-Seq, stLFR, and 10x linked-reads from ATCC-MSA-1003.

**Supplementary Table 8**

The genome fractions generated by different assembly tools for the five strains with abundances of 0.02% using TELL-Seq, stLFR, and 10x linked-reads from ATCC-MSA-1003.

**Supplementary Table 10**

The genome fractions, per-strain NA50 and NGA50 generated by different assembly tools for the 10 strains using the stLFR linked-reads from ZYMO.

**Supplementary Table 11**

The statistics of MAGs generated for the three human gut microbiomes by all assemblers.

**Supplementary Table 12**

The closest reference genomes of the NCMAGs with species-level annotations generated by all assemblers and their alignment statistics reported by GTDB-Tk (v2.1.0).

**Supplementary Table 13**

The genome fractions, total assembly length for contigs longer than 10Kb, per-strain NA50 and NGA50 by different assemblers using short- and long-reads from the two mock communities.

