## Supplementary Text and Figures for "Exploring high-quality microbial genomes by assembling short-reads with long-range connectivity"

### Supplementary Notes

#### Supplementary Note 1: 10x, stLFR and TELL-Seq Sequencing

We compared the characteristics of 10x, stLFR and TELL-Seq linked-reads on the mock community. 10x linked-reads generated the lowest number of unique barcodes (2.31 million; **Table 1.1**), where the values of stLFR and TELL-Seq were 45.38 and 16.61 million, respectively (**Table 1.1**). We reconstructed the physical long fragments and calculated the number of fragments per barcode ($N_{F/B}$) for the three technologies by mapping the linked-reads to reference genomes. 10x linked-reads had the lowest barcode specificity with $N_{F/B}$ of 16.61 (**Table 1.1**), where the values of stLFR and TELL-Seq linked-reads were much lower, which were 1.54 and 4.26, respectively (**Table 1.1**). We aggregated the barcodes based on long fragments they involved and found highest fraction is 6.41% for 14 long fragments. The highest fractions for both stLFR (62.98%) and TELL-Seq (27.05%) linked-reads were matching to one long fragment (**Figure 1.1**).

10x and stLFR linked-reads generated comparable lengths of the long fragments, that were longer than those from TELL-Seq linked-reads (**Figure 1.2**); the weighted average of long fragment lengths ($W_{\mu_{FL}}$) for 10x, stLFR and TELL-Seq linked-reads were 17.02Kb, 15.68Kb and 11.85Kb, respectively (**Table 1.1**). All the three linked-read sequencing platforms had shallow depth of short-reads for long fragments, with similar distributions (**Figure 1.3**) and comparable average value ($C_R$: 10x = 0.21, stLFR = 0.20, TELL-Seq = 0.17; **Table 1.1**).

Further, we investigated the insert sizes of linked-reads generated from the three sequencing platforms. stLFR (246 bp) and TELL-Seq (204 bp) linked-reads obtained lower median insert sizes than 10x linked-reads (339 bp). We observed the histograms of the insert sizes from stLFR and TELL-Seq linked-reads were more biased than that from 10x linked-reads, and the curve of histogram from TELL-Seq linked-reads had frequent vibrations (**Figure 1.4**).

| | 10x linked-reads | stLFR linked-reads | TELL-Seq linked-reads |
|---|---|---|---|
| Number of barcodes | 2,305,295 | 45,381,261 | 16,609,060 |
| Number of fragments per barcode ($N_{F/B}$) | 16.61 | 1.54 | 4.26 |
| Weighted fragment length ($W_{\mu_{FL}}$, bp) | 17,022 | 15,683 | 11,848 |
| Read depth per fragment ($C_R$) | 0.21 | 0.20 | 0.17 |
| Median insert size (bp) | 339 | 246 | 204 |

**Table 1.1.** The statistics of 10x, TELL-Seq and stLFR linked-reads.

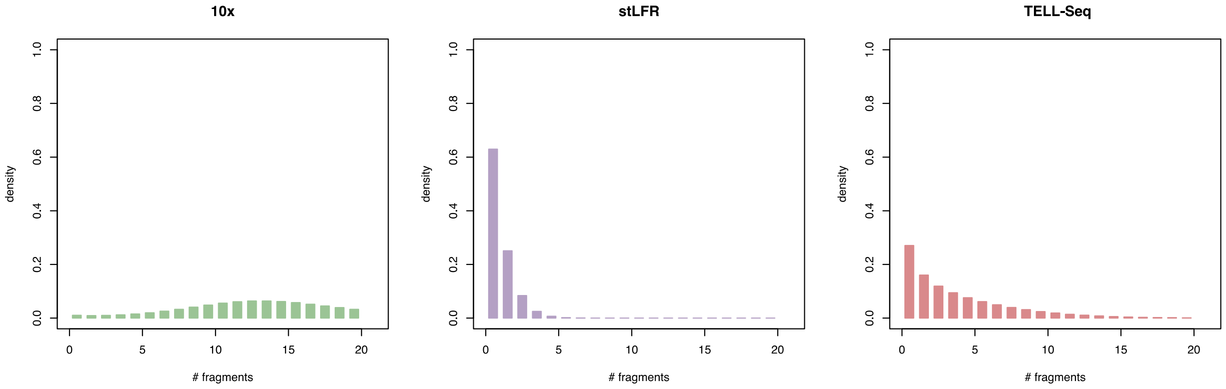

**Figure 1.1.** The distributions of the number of physical long fragments per barcode.

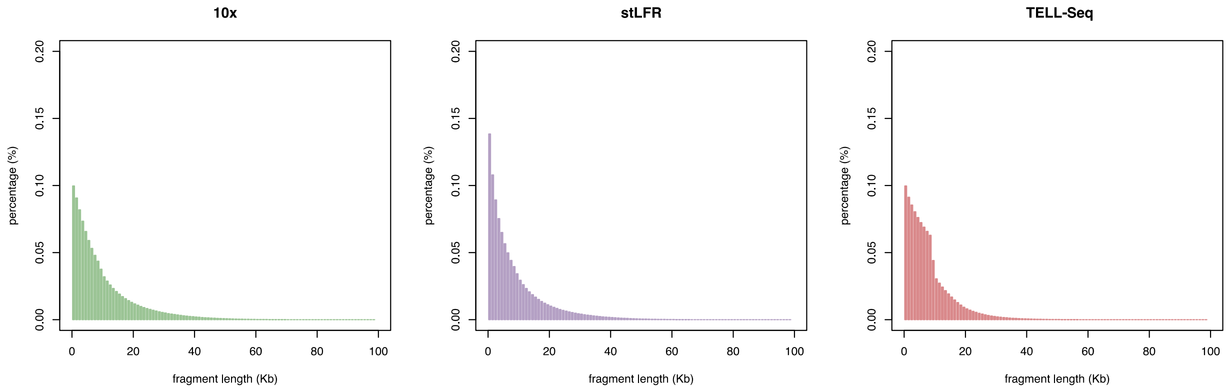

**Figure 1.2.** The distributions of the lengths of physical long fragments.

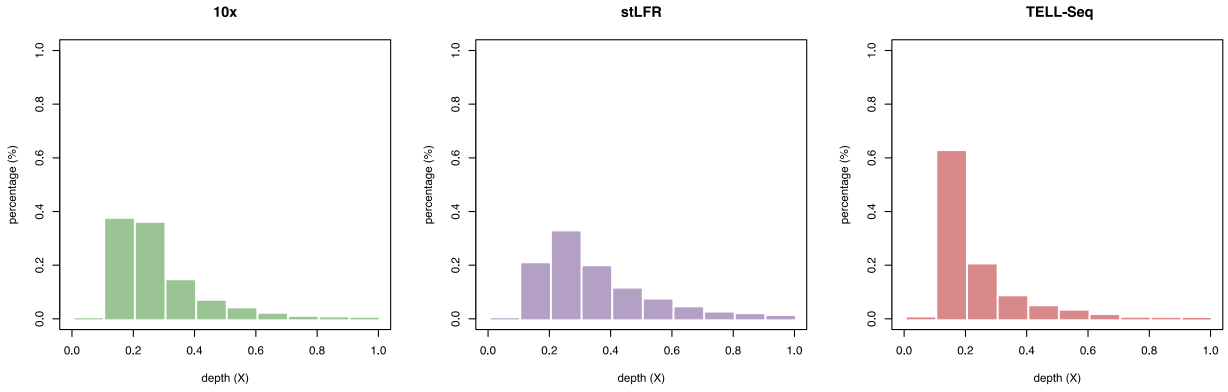

**Figure 1.3.** The distributions of read depths per long fragments.

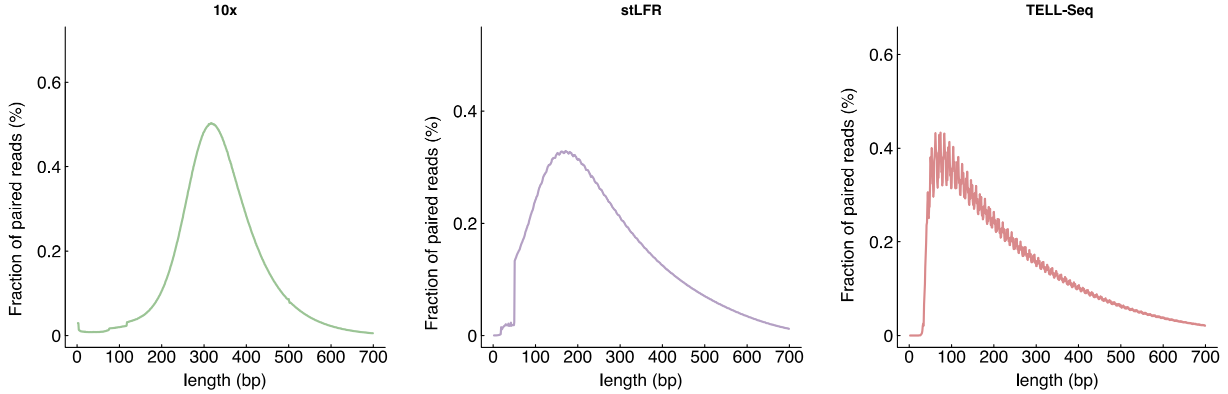

**Figure 1.4.** The distributions of insert sizes for 10x, stLFR and TELL-Seq linked-read.

#### Supplementary Note 2: The architecture of VAE

We used VAE to learn the low-dimensional representation of the co-barcoded linked-reads based on their abundances ($X_A$) and TNF ($X_T$) features (**Figure 2.1**). The input of VAE was the concatenation of L1-normalized $X_A$ (400 dimensions) and $X_T$ (136 dimensions). In the encoder, the input features were transformed by two fully connected hidden layers with 512 neurons for each, and compressed into two 32-dimensional latent parameters $\mu$ and $\sigma$ for Gaussian distribution. We sampled the latent embedding $Z$ from the Gaussian distribution $N(\mu, \sigma^2)$. To make the sampling process trainable for $\mu$ and $\sigma$, we used the reparameterization trick, that calculated $Z$ by $Z = \mu + \sigma \circ \epsilon, \epsilon \sim N(0, 1)$. In the decoder, we used two fully connected hidden layers with the same settings as the encoder, and reconstructed the input features using an output layer with 516 neurons. The output vector was split to the reconstructed abundances ($\hat{X}_A$) and TNF ($\hat{X}_T$) features according to their corresponding dimensions. Both $\hat{X}_A$ and $\hat{X}_T$ were transformed by softmax before output. To increase the stability of training, each hidden layer in the encoder and decoder was followed by a batch normalization layer and a dropout layer with P=0.2. Our batch size and learning rate were set to 2,048 and 0.005. We exploited weight decay of 1e-4 by default to avoid overfitting and used the early stopping that stopped the training when the loss started to increase, which substantially reduced the training time.

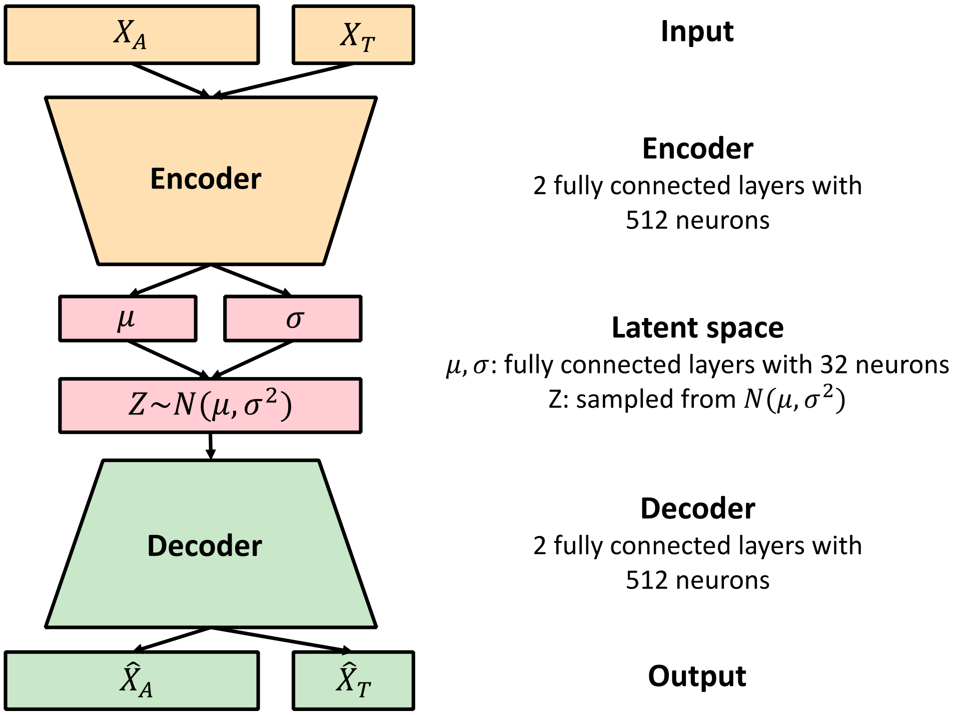

**Figure 2.1.** The architecture of VAE used in Pangaea.

#### Supplementary Note 3: Parameter search of Pangaea

##### 1. Determining $\alpha$ and $\beta$ for the weights in the loss function

We searched a range of hyper-parameters $\alpha$ and $\beta$ on the stLFR linked-reads dataset from ATCC-MSA-1003. The overall adjusted rand index (ARI) and F1 did not vary much when changing $\beta$, so we used $\beta$ =0.015 because the opposite trends in overall precision and ARI were observed after $\beta$ =0.015 (**Figure 3.1 a**). The lowest $\alpha$ resulted in the highest overall precision, recall, F1, and ARI (**Figure 3.1 b**). Therefore, we adopted a low value of $\alpha$ =0.1. The $w_T$ is much larger than $w_A$, which means TNF is a more important feature than the *k*-mer histogram in read binning. This might be because the *k*-mer-histograms using a larger *k* (*k*=15) than TNF (*k*=4) are more likely to be influenced by sequencing errors and contain more noise. We used a small $w_{KL}$, which was consistent with the previous study that applies VAE in contig binning tasks [1].

##### 2. Determining the range of thresholding T for multi-thresholding reassembly

We investigated the impact of the range of thresholds T on the final assembly results using the TELL-Seq dataset of ATCC-MSA-1003 (**Supplementary Table 5**). We compared the results without multi-thresholding reassembly (i.e., T = {}), with T = {10}, T = {10, 30}, T = {10, 30, 50}, T = {10, 30, 50, 70} and T = {10, 30, 50, 70, 90}. We observed T = {10, 30} obtained the highest N50 (1.36MB) and the second higher overall NA50 (649.67Kb; only slightly lower than the overall NA50 produced by T = {10, 30, 50, 70}; **Supplementary Table 5**). Involving higher thresholds did not bring much improvement and would need a longer time for running (**Supplementary Table 5**). Therefore, we unified the default range of thresholds as T = {10, 30}.

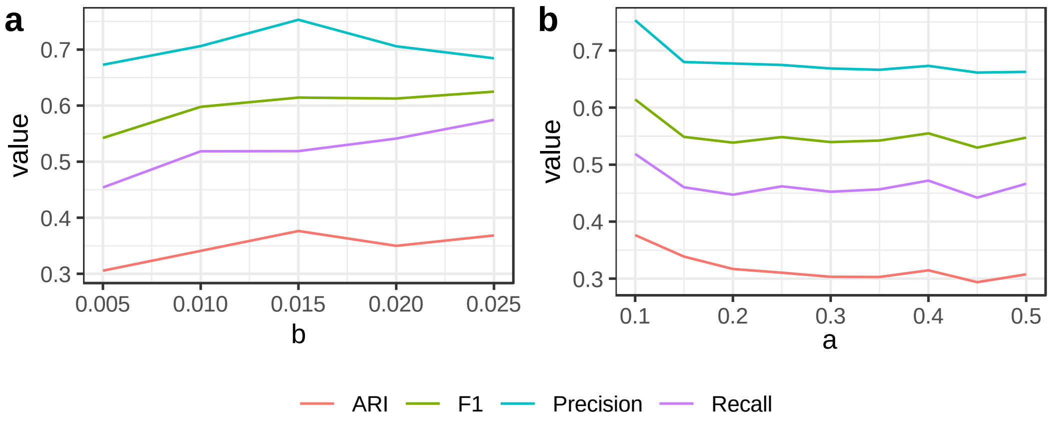

**Figure 3.1.** The binning performance on the stLFR linked-reads dataset of ATCC-MSA-1003 using different $\alpha$ (subfigure **a**, $\beta$ is fixed at 0.015) and $\beta$ (subfigure **b**, $\alpha$ is fixed at 0.1) in the loss function of VAE.

#### Supplementary Note 4: Pangaea generated more NCMAGs than PacBio long-read sequencing

We compared the contigs from Pangaea with those from metaFlye on PacBio CLR long-reads of S1 and S2 (**Supplementary Table 3**; **Supplementary Figure 4**). Although metaFlye generated contigs with higher N50s, Pangaea produced a substantially greater total assembly length for both S1 (Pangaea = 488.19Mb, metaFlye = 243.88Mb) and S2 (Pangaea = 408.82Mb, metaFlye = 256.78Mb; **Table 4.1**). Moreover, Pangaea generated significantly more NCMAGs than metaFlye (Pangaea = 41, metaFlye = 16; **Figure 4.1 a**), especially those with N50s smaller than 1Mb (Pangaea = 29, metaFlye = 4; **Figure 4.1 b and d**) and read depths lower than 300x (Pangaea = 27, metaFlye = 0; **Figure 4.1 c**), whereas Pangaea and metaFlye obtained comparable numbers of NCMAGs with N50 larger than 1Mb (Pangaea = 12, metaFlye = 12; **Figure 4.1 b**). Pangaea also assembled more ORF clusters (Pangaea = 522.81K, metaFlye = 169.79K), 5s rRNAs (Pangaea = 612, metaFlye = 400), 16s rRNAs (Pangaea = 690, metaFlye = 492), 23s rRNAs (Pangaea = 700, metaFlye = 491), plasmid contigs (Pangaea = 3,376, metaFlye = 218) and viral contigs (Pangaea = 1,707, metaFlye = 61) than metaFlye (**Table 4.2**).

| | Human gut microbiome (S1) | | Human gut microbiome (S2) | |
|---|---|---|---|---|
| | Pangaea | metaFlye | Pangaea | metaFlye |
| Total assembly length | 488,785,611 | 243,883,392 | 408,819,148 | 256,776,331 |
| Largest contig | 2,394,379 | 3,388,254 | 2,877,256 | 4,327,156 |
| N50 | 64,394 | 168,442 | 188,161 | 239,008 |

**Table 4.1.** Assembly statistics for Pangaea and metaFlye on S1 and S2.

| | Pangaea | metaFlye |
|---|---|---|
| ORF clusters | 522,812 | 169,785 |
| 5S rRNA | 612 | 440 |
| 16s rRNA | 690 | 492 |
| 23s rRNA | 700 | 491 |
| Plasmid contigs (>1Kb) | 3,376 | 218 |
| Viral contigs (>1Kb) | 1,707 | 61 |

**Table 4.2.** The total number of ORF clusters, rRNAs, plasmid contigs (>1Kb), and viral contigs (>1Kb) of Pangaea and metaFlye on S1 and S2.

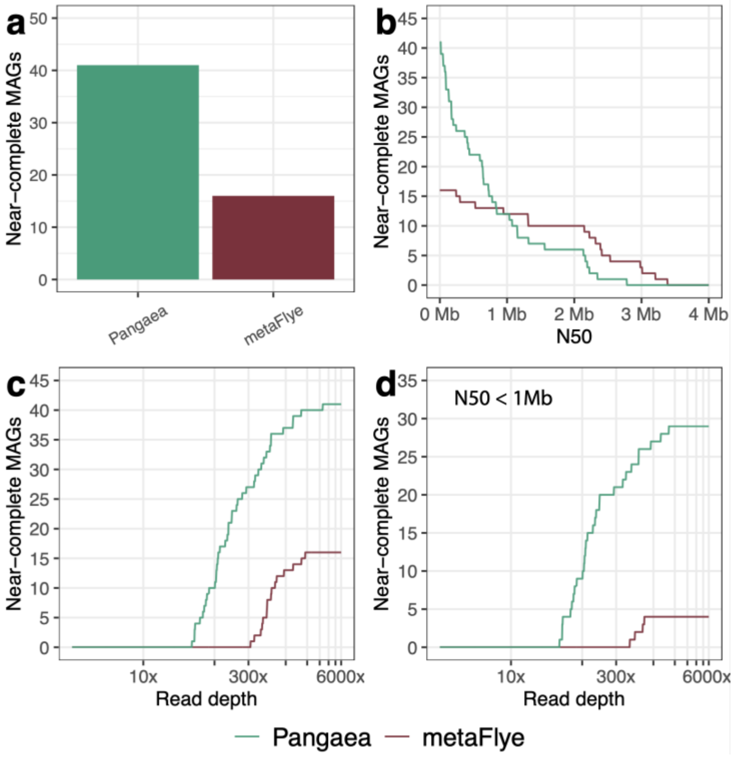

**Figure 4.1.** The number of NCMAGs of Pangaea and metaFlye on S1 and S2 (a), the NCMAGs of Pangaea and metaFlye on S1 and S2 with a minimum value of N50 (b), the NCMAGs of Pangaea and metaFlye on S1 and S2 with a maximum value of read depth (c), and the NCMAGs (N50 < 1Mb) of Pangaea and metaFlye on S1 and S2 with a maximum value of read depth (d).

### Supplementary Figures

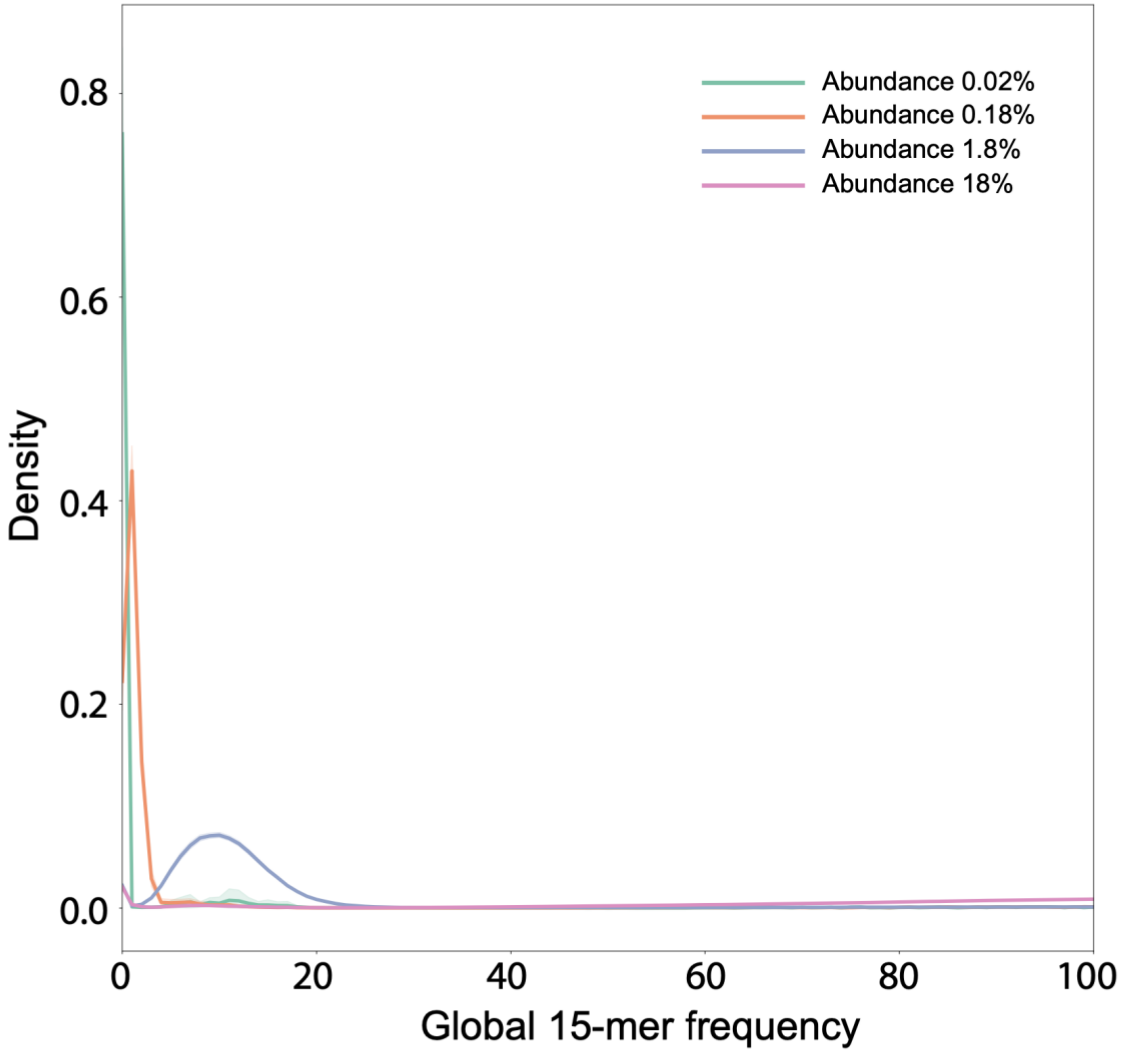

**Supplementary Figure 1.** The *k*-mer histograms of co-barcoded reads of different abundance levels on the stLFR linked-reads of ATCC-MSA-1003. The line for abundance 18% is truncated at the global 15-mer frequency at 100 for better visualization of the other three abundance levels.

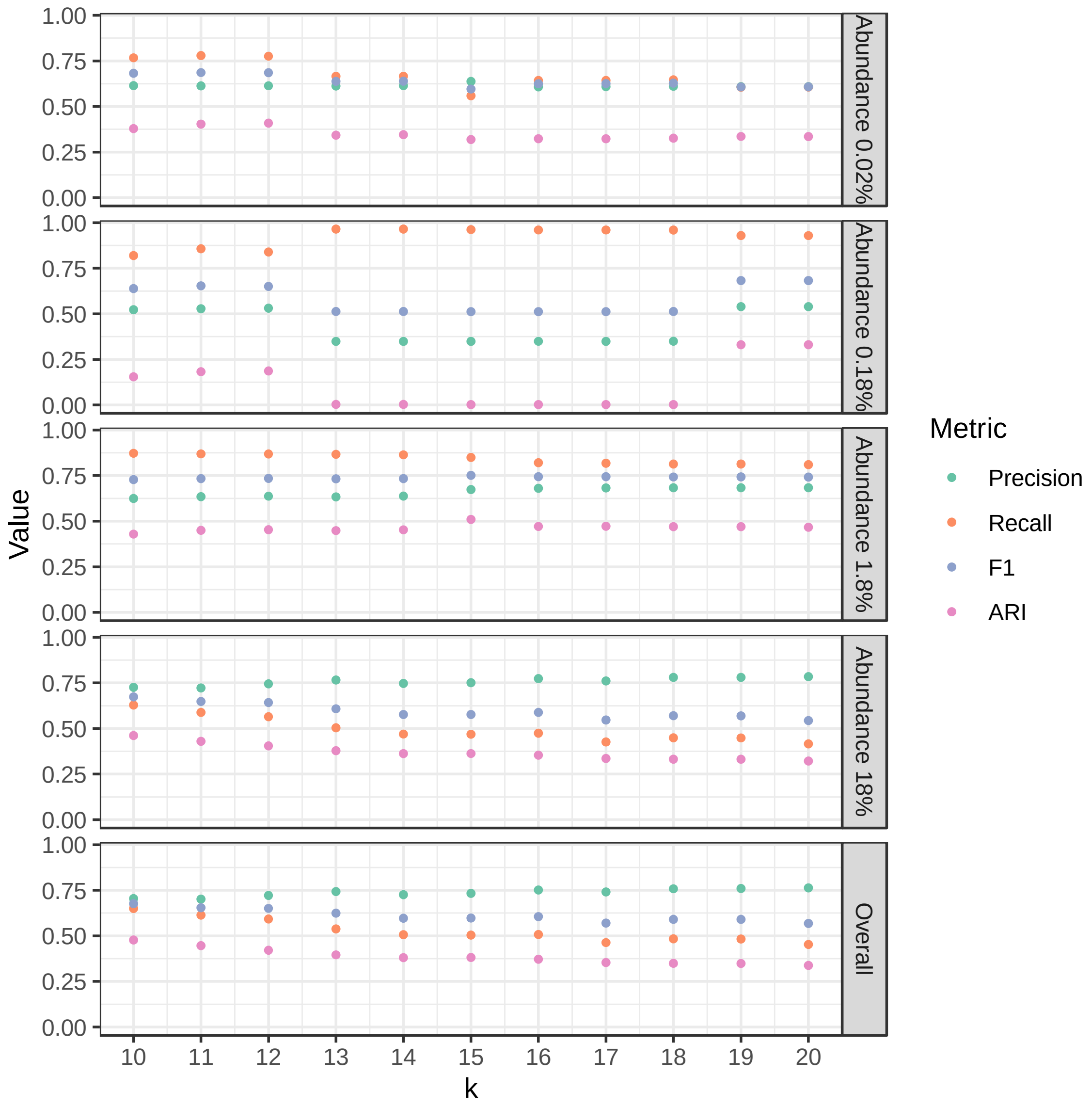

**Supplementary Figure 2.** The precision, recall, F1 score and adjusted rand index (ARI) values of the barcode binning results using different numbers of clusters on the stLFR linked-reads of ATCC-MSA-1003.

**a**

| | Pangaea (k=10) | Pangaea (k=15) | Pangaea (k=20) |
|---|---|---|---|
| # contigs | 3585 | 3594 | 3609 |
| Largest contig | 4972897 | 3348227 | 4972369 |
| Total length | 58494697 | 58259182 | 58311142 |
| N50 | 1987506 | 1833445 | 1833833 |
| N75 | 373579 | 396900 | 373473 |
| L50 | 10 | 12 | 11 |
| L75 | 26 | 27 | 27 |
| Genome fraction (%) | 84.873 | 85.045 | 82.155 |
| NA50 | 679104 | 732394 | 746839 |
| NA75 | 206301 | 223106 | 195429 |
| LA50 | 25 | 24 | 22 |
| LA75 | 61 | 57 | 56 |

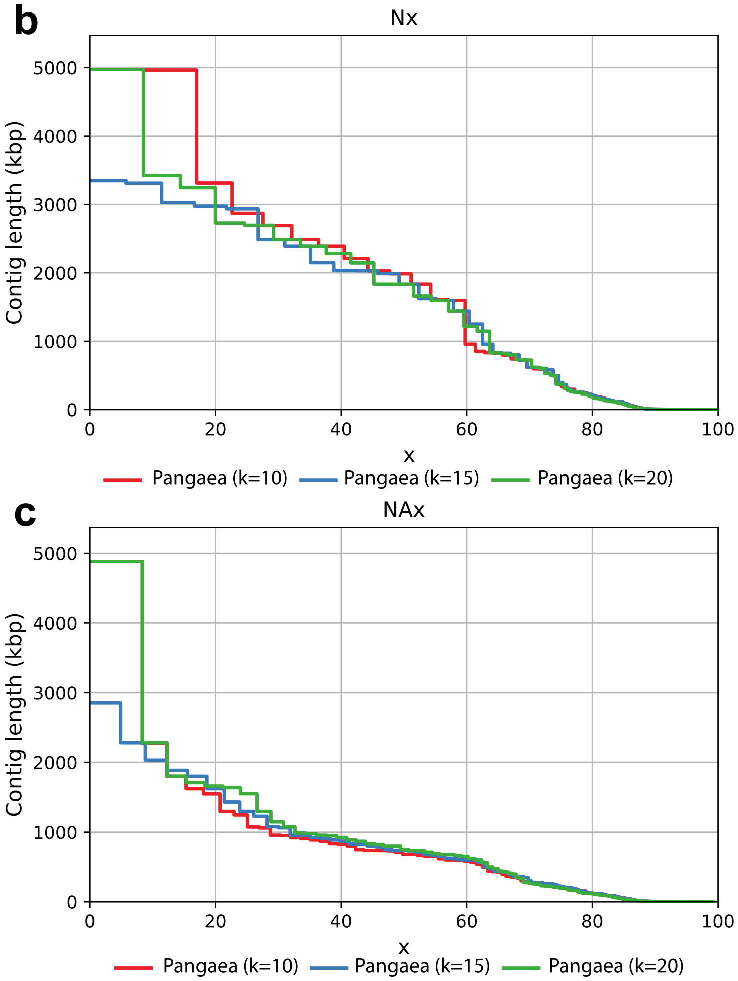

**Supplementary Figure 3.** The assembly statistics (**a**), Nx (**b**), and NAx (**c**) of Pangaea assemblies on the stLFR linked-reads of ATCC-MSA-1003 based on different cluster numbers (k=10, 15, and 20).

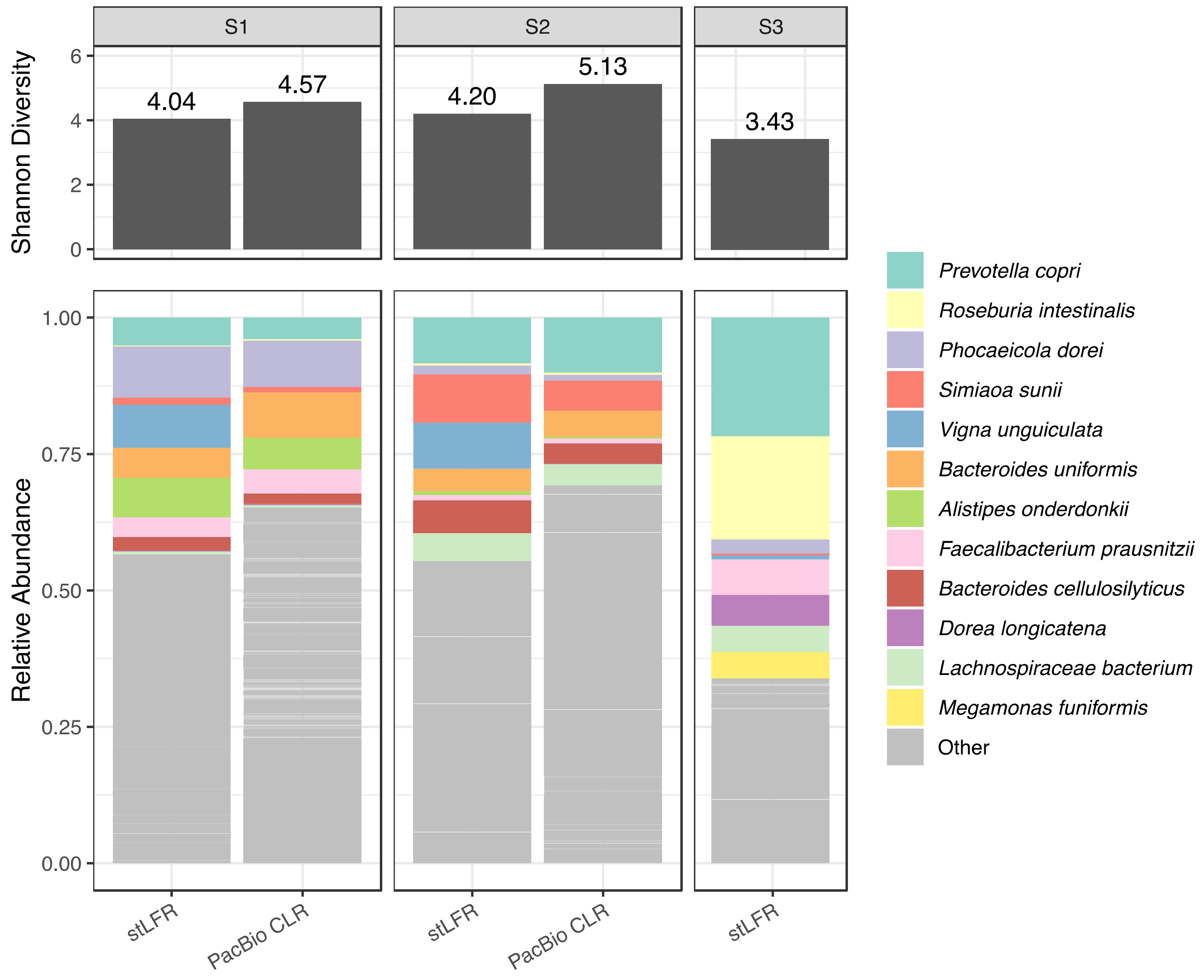

**Supplementary Figure 4.** The reads composition and Shannon diversity of stLFR linked-reads and PacBio CLR long-reads on human gut microbiomes.

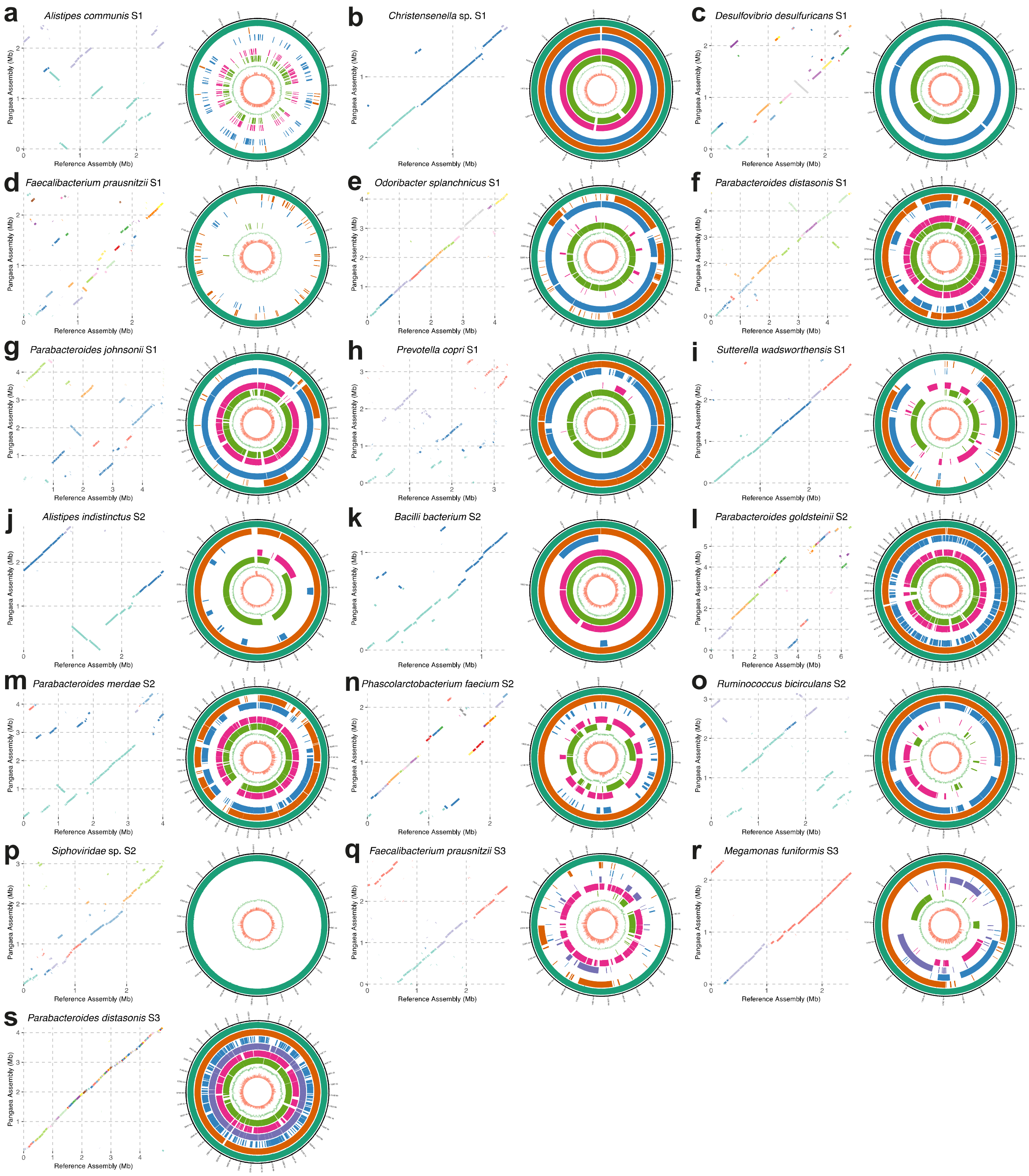

**Supplementary Figure 5**. Genome collinearity analysis between the selected NCMAGs produced by Pangaea and their closest reference genomes (dot plots), and comparison of the corresponding MAGs produced by different assemblers (circos plots) from S1, S2 and S3. Colors are used in the dot plots to distinguish different contigs. The eight rings in the circos plots from outside to inside denote the Pangaea MAGs (dark green), Athena MAGs (orange), Supernova MAGs (blue), cloudSPAdes MAGs (purple), MEGAHIT MAGs (pink), metaSPAdes MAGs (light green), GC-skew of Pangaea MAGs, and read depth of Pangaea MAGs, respectively. If the same species was annotated by more than one MAG from the same assembler, the one with the highest N50 is shown here.

### Supplementary Tables

| Organism | ATCC ID | Composition | Taxonomy ID |
| --- | --- | --- | --- |
| *Bacteroides vulgatus* | ATCC_8482 | 0.02% | NCBI:txid435590 |
| *Bifidobacterium adolescentis* | ATCC_15703 | 0.02% | NCBI:txid367928 |
| *Deinococcus radiodurans* | ATCC_BAA816 | 0.02% | NCBI:txid243230 |
| *Enterococcus faecalis* | ATCC_47077 | 0.02% | NCBI:txid474186 |
| *Schaalia odontolytica* | ATCC_17982 | 0.02% | NCBI:txid411466 |
| *Acinetobacter baumannii* | ATCC_17978 | 0.18% | NCBI:txid400667 |
| *Cutibacterium acnes* | ATCC_11828 | 0.18% | NCBI:txid1091045 |
| *Helicobacter pylori* | ATCC_700392 | 0.18% | NCBI:txid85962 |
| *Lactobacillus gasseri* | ATCC_33323 | 0.18% | NCBI:txid324831 |
| *Neisseria meningitidis* | ATCC_BAA335 | 0.18% | NCBI:txid122586 |
| *Bacillus cereus* | ATCC_10987 | 1.8% | NCBI:txid222523 |
| *Clostridium beijerinckii* | ATCC_35702 | 1.8% | NCBI:txid864803 |
| *Pseudomonas aeruginosa* | ATCC_9027 | 1.8% | NCBI:txid287 |
| *Staphylococcus aureus* | ATCC_BAA1556 | 1.8% | NCBI:txid451515 |
| *Streptococcus agalactiae* | ATCC_BAA611 | 1.8% | NCBI:txid208435 |
| *Escherichia coli* | ATCC_700926 | 18% | NCBI:txid511145 |
| *Porphyromonas gingivalis* | ATCC_33277 | 18% | NCBI:txid431947 |
| *Rhodobacter sphaeroides* | ATCC_17029 | 18% | NCBI:txid349101 |
| *Staphylococcus epidermidis* | ATCC_12228 | 18% | NCBI:txid176280 |
| *Streptococcus mutans* | ATCC_700610 | 18% | NCBI:txid210007 |

**Supplementary Table 1.** The composition and taxonomy ID of the 20 strains in the ATCC-MSA-1003 mock community.

| Microbial community | Sequencing technology | Read length | Total sequencing size (Gb) |
|---|---|---|---|
| ZYMO | stLFR | 100 | 143.04 |
| ZYMO | ONT | 4,504 (average) | 2.48 (downsampled) |
| ATCC-MSA-1003 | 10x | 150 | 100.38 |
| ATCC-MSA-1003 | TELL-Seq | 146 | 173.28 |
| ATCC-MSA-1003 | stLFR | 100 | 132.95 |
| ATCC-MSA-1003 | PacBio CLR | 8,875 (average) | 1.27 (downsampled) |
| Human gut microbiome (S1) | stLFR | 100 | 136.60 |
| Human gut microbiome (S1) | PacBio CLR | 8,878 (average) | 6.26 |
| Human gut microbiome (S2) | stLFR | 100 | 131.59 |
| Human gut microbiome (S2) | PacBio CLR | 8,973 (average) | 8.39 |
| Human gut microbiome (S3) | stLFR | 100 | 50.74 |

**Supplementary Table 2.** The sequencing statistics of the datasets.

| | Binning of Pangaea | METABCC-LR |
|---|---|---|
| Number of clusters | 15 | 2 |
| F1 – abundance 0.02% | 0.6073 | 0.4624 |
| ARI – abundance 0.02% | 0.3246 | 0 |
| F1 – abundance 0.18% | 0.5125 | 0.5167 |
| ARI – abundance 0.18% | 0.0022 | 0.0140 |
| F1 – abundance 1.8% | 0.7500 | 0.6155 |
| ARI – abundance 1.8% | 0.5069 | 0.1740 |
| F1 – abundance 18% | 0.5960 | 0.6351 |
| ARI – abundance 18% | 0.3581 | 0.2019 |
| F1 - overall | 0.6144 | 0.5887 |
| ARI - overall | 0.3764 | 0.1704 |

**Supplementary Table 3.** The F1 scores and ARIs of the binning algorithm of Pangaea and METABCC-LR on the stLFR linked-reads of ATCC-MSA-1003.

| | Pangaea TELL-Seq | Pangaea stLFR | Athena 10x | Supernova 10x | cloudSPAdes 10x |
|---|---|---|---|---|---|
| Total assembly length | 61,990,266 | 59,485,233 | 52,159,979 | 89,828,047 | - |
| Genome fraction (%) | 82.63 | 84.43 | 77.20 | 75.08 | - |
| Longest alignment | 4,968,123 | 2,853,278 | 2,278,020 | 974,529 | - |
| Overall N50 | 1,360,322 | 1,619,916 | 596,076 | 32,128 | - |
| Overall NA50 | 649,672 | 731,990 | 437,889 | 30,194 | - |
| NGA50 per strain | 887,107 | 677,353 | 338,609 | 89,994 | - |
| NA50 per strain | 838,457 | 628,059 | 337,373 | 93,098 | - |

**Supplementary Table 6.** Assembly statistics for different linked-read sequencing platforms on the ATCC-MSA-1003 mock community. cloudSPAdes of 10x linked-reads was unavailable because it requires large memory (>1TB) on the dataset.

| Organism | Composition |
| --- | --- |
| *Listeria monocytogenes* | 89.1% |
| *Pseudomonas aeruginosa* | 8.9% |
| *Bacillus subtilis* | 0.89% |
| *Saccharomyces cerevisiae* | 0.89% |
| *Escherichia coli* | 0.089% |
| *Salmonella enterica* | 0.089% |
| *Lactobacillus fermentum* | 0.0089% |
| *Enterococcus faecalis* | 0.00089% |
| *Cryptococcus neoformans* | 0.00089% |
| *Staphylococcus aureus* | 0.000089% |

**Supplementary Table 9.** The composition of the 10 strains in the ZYMO mock community.
